# Multiscale computational framework for generating vascularized biohybrid tissue constructs

**DOI:** 10.64898/2026.02.28.708633

**Authors:** Andrew A. Guy, Alexander W. Justin, Athina E. Markaki

## Abstract

Vascularization remains a central challenge in building large-scale biohybrid tissues that integrate living and synthetic components. Without a perfusable vascular network, nutrient delivery and waste removal become insufficient, leading to hypoxia and a loss of viability in thicker tissue constructs. We present Lattice Sequence Vascularization (LSV), a multiscale computational design framework for generating hierarchical, biomimetic vascular networks that are compatible with 3D-printing constraints and manufacturable within arbitrary geometries. LSV employs a divide-and-conquer strategy in which vessels grow and remodel at a specified terminal scale before recursively subdividing to form the full hierarchy. By enforcing hierarchy, LSV produces networks that exhibit self-similarity across length scales, a defining feature of physiological vasculature. The framework integrates synthetic considerations (e.g., hydrogel permeability), biological constraints (Murray’s law, cross-scale biomimicry, organ-specific perfusion requirements) and manufacturing requirements relevant to 3D printing and microfabrication. We demonstrate the incorporation of capillary-scale functional substructures (e.g. organoid traps) and the generation of complex architectures with multiple inlets and outlets (e.g. liver-like geometries), enabling organ-scale vasculature design.

## 1 Introduction

Generating tissue-mimetic vasculature is a key challenge for tissue engineering and 3D cell culture systems [1, 2]. Biohybrid materials, which integrate living biological components with engineered synthetic scaffolds, offer a versatile platform for constructing complex tissue-like constructs with pre-patterned vasculature. Within native tissue, oxygen and nutrients are efficiently delivered via a bifurcating channel network comprising arteries and veins, which feed a highly interconnected capillary bed, capable of constant remodelling to meet changing oxygen and nutrient requirements. Branching vessels, which distribute flow from a single inlet to multiple terminals, allow for better coverage of a 3D space, while also homogenizing the pressure drop from flow across any unit volume of tissue. Further, the complexity of vasculature found in living tissue is the result of a continuous optimisation of metabolite requirements during the growth and development of the organism.

A wide range of biofabrication techniques have been developed to generate functional vasculature in biohybrid materials [3]. Microfluidic devices commonly incorporate single uniaxial channels [4]; however, these pre-patterned channels are typically positioned around the periphery of the tissue construct, limiting its size in all dimensions. Sacrificial bioprinting and stereolithography have enabled fabrication of channel networks in 3D [5–7], but these channels often consist of simple uniaxial arrays that are not optimised for effective nutrient and oxygen supply into the surrounding tissue space. Although vasculature can form biologically through cellular self-organisation and angiogenesis [8, 9], this process is confined to the capillary scale, resulting in highly interconnected networks without the hierarchical branching of larger vessels.

The design, simulation and fabrication of vasculature with native anatomical complexity present a significant challenge [10, 11]. Designing even simple bifurcating networks manually within computer-aided design (CAD) packages is a time intensive exercise, and vasculature of the level of complexity found in native tissue is unfeasible this way, and so techniques are required to procedurally generate such vascular networks to support any desired space, subject to user-defined physiological parameters.

Several algorithms have been developed to generate arterial trees, accounting for the hierarchical organization of vasculature and the physiological constraints of blood flow. Among these, Constrained Constructive Optimization (CCO) and Space Colonization (SC) are iterative growth methods, which progressively expand the network by perfusing new regions of space. CCO was initially developed to model single arterial trees in 2D [12], and later extended to 3D [13]. The concept of “staged growth” was introducted in subsequent work [14] to enforce a more balanced hierachical structure. CCO has proven effective in generating large, biomimetic vascular networks with customizable terminal flow rates and uniform pressures. However, it has a number of deficits such as high bifurcation asymmetry and an inability to handle non-convex domains, requiring either a predefined growth staging sequence or a computationally expensive penalty function augmentation. The Adaptive-CCO [15] introduces staged growth, where terminal sites develop in phases based on probability distributions, allowing for early hierarchical structure formation and gradual refinement, unlike the original CCO, which tends to generate bifurcations immediately. In SC, networks grow outward from the root towards a set of “auxin sources”, mimicking biological processes such as nutrient-driven flow [16, 17]. However, unlike CCO, SC does not ensure complete perfusion (by virtue of allowing terminal nodes to become bifurcations) and lacks optimization for any network-scale physiological cost function, potentially limiting its effectiveness in artificial designs.

Approaches, such as Global Constructive Optimization (GCO), Simulated Annealing (SA), and Graph-Based Methods, fall under the category of iterative rearrangement. GCO generates vascular networks by initially connecting all terminals and refining connections [18]. While it enables higher-order branching, which CCO lacks, it assumes flow distributions that may diverge from physiological conditions [19]. SA is well-suited for non-convex domains, but it is computationally expensive for large or multiple vascular networks due to the vast number of possible configurations [20]. Graph-Based Methods constrain node and edge placement to a subset of a predefined graph, making it challenging to generate realistic or physiologically optimal networks. These methods may also result in “floating leaf vessels”, hindering both fabrication and perfusion. Additionally, they are computationally expensive, especially in 3D domains [21]. Importantly, GCO, SA and Graph-based methods do not inherently guarantee uniform perfusion.

Methods such as fractals [22] are predetermined, while Lindenmayer Systems [23] are stochastically generated according to a rigid set of rules. As a result neither method accounts for the evolving state of the network during iteration. In addition, they make it difficult to guarantee non-overlapping vessels or adequate perfusion [24]. From a tissue engineering perspective, CCO and its adaptations are the preferred approaches for generating large, biomimetic networks, as they allow for arbitrary specification of terminal flow rates and uniform terminal pressures. Further, GCO and SA offer topological optimisation capabilities that can be integrated into the design process.

Most early work focused on modelling single arterial trees, typically involving only one network and rarely more than two. Initial efforts to model dual networks were carried out by [25–27] with subsequent refinements incorporating the Fåhræus-Lindqvist effect [28, 29] at the smallest scales. However, these methods do not scale well. Their optimization routines require constant intersection checks, resulting in suboptimal networks generated at substantial computational costs with poor scaling. Moreover, they fall short in guaranteeing adequate perfusion, because there may not be a valid connection available for some terminal sites. We previously introduced the Accelerated-CCO (A-CCO) algorithm [30], the first vascular design framework to support an arbitrary number of interpenetrating networks, a scheme for collision resolution between multiple trees and the first to be compatible with 3D printing constraints. We demonstrated that computational complexity can be significantly reduced by exploiting the tree structure and only evaluating candidate paths whilst they continue making progress towards the target, a strategy that has since been independently validated by [31]. This study [31] featured a “hybrid” topological optimization approach, in which SA was used as a dedicated topological optimization step between CCO iterations. Physiological constraints including Murray’s law and prescribed terminal flow rates were treated as constraints, and the Fåhræus-Lindqvist effect was accounted for. Building upon the mathematical optimization techniques introduced in [31], the framework was extended to non-convex domains and multiple networks such as the brain gyrus [32] and liver [33], without relying on CCO. Recently, building on Schreiner and Karch’s original CCO framework [12, 13], an open-source CCO algorithm (svVascularize) [34] was used to generate vascular networks in complex geometries, which were subsequently 3D printed as perfusable models. The networks were optimised for minimum vascular volume, consistent with earlier CCO work [12, 25, 31, 35], and flow behaviour was validated using 0D and 1D models together with computational fluid dynamics (CFD) simulations of a planar network [34].

The generation of clinically relevant biohybrid tissue constructs requires algorithmic strategies capable of producing vascular architectures that support uniform oxygen and nutrient transport throughout the tissue volume. Such networks should be optimisable with respect to a range of efficiency-type cost functions, including configurations with fixed inlet and outlet radii for coupling to external perfusion systems, as commonly employed in tissue engineering (e.g. [34, 36, 37]).

Here, we present a computational framework that autonomously generates vascular networks optimised for metabolic costs, thereby enabling the design of energetically efficient constructs. The framework can navigate geometric bottlenecks, hierarchically generate major vessels, and ensure comprehensive terminal scale perfusion throughout the biohybrid construct. To enhance biomimetic fidelity across length-scales, the approach incorporates adaptive physiological constraints, including a variable implementation of Murray’s law, and supports the integration of functional microarchitectural features within engineered tissues, such as organoid trapping structures. Algorithmic performance is systematically evaluated across representative domains, demonstrating both scalability and robustness.

In contrast to existing methodologies that rely on random sampling or computationally expensive coupled oxygen transport, the proposed algorithm avoids stochastic sampling entirely. Instead, it is based on two key assumptions: a critical length governing cell-channel distance and a uniform perfusion criterion, with terminal nodes placed on lattices to ensure computational efficiency and reliability. Terminal nodes, initially placed on lattices, are permitted to move within a prescribed neighbourhood, allowing modest cost improvements whilst still satisfying the critical length constraint. Across a broad range of domain geometries, this formulation guarantees that the cell–channel distance requirement is satisfied or, where infeasible, provides immediate and unambiguous identification of incomplete growth.

The present work begins with exposition of the key assumptions underlying our approach: (a) a suitable starting point for designing feasible vasculature for tissue engineering is to ensure that every point within the domain lies within a critical length 𝓁 of a terminal channel; and (b) that certain assumptions can be made to enable rapid computation of branch/node properties without sacrificing uniform perfusion.

Building on these assumptions, we introduce a novel algorithm, Lattice Sequence Vascularization (LSV), with 𝒪 (*N* log *N*) complexity, where *N* ≈ |Ω|/𝓁^3^ is the number of ‘unit cells’ for a given terminal spacing and tissue domain Ω, which ensures that the critical distance constraint is met subject to fairly lax constraints. Our framework handles inter- and intra-network collision resolution while accounting for mesh boundaries. In addition, it supports optimization for a range of costs depending on the intended function. We demonstrate how the framework can be adapted to improve biomimicry and enforce physiological constraints. This work highlights the broad design possibilities made feasible by the developed software libraries. High-level descriptions of the algorithms are provided in the appendices (with full details available in a reference implementation), while suitable parameter choices depend heavily on manufacturing requirements. An overview of the software workflow, including relevant inputs and outputs, is illustrated in Figure 1.

**Figure 1:**
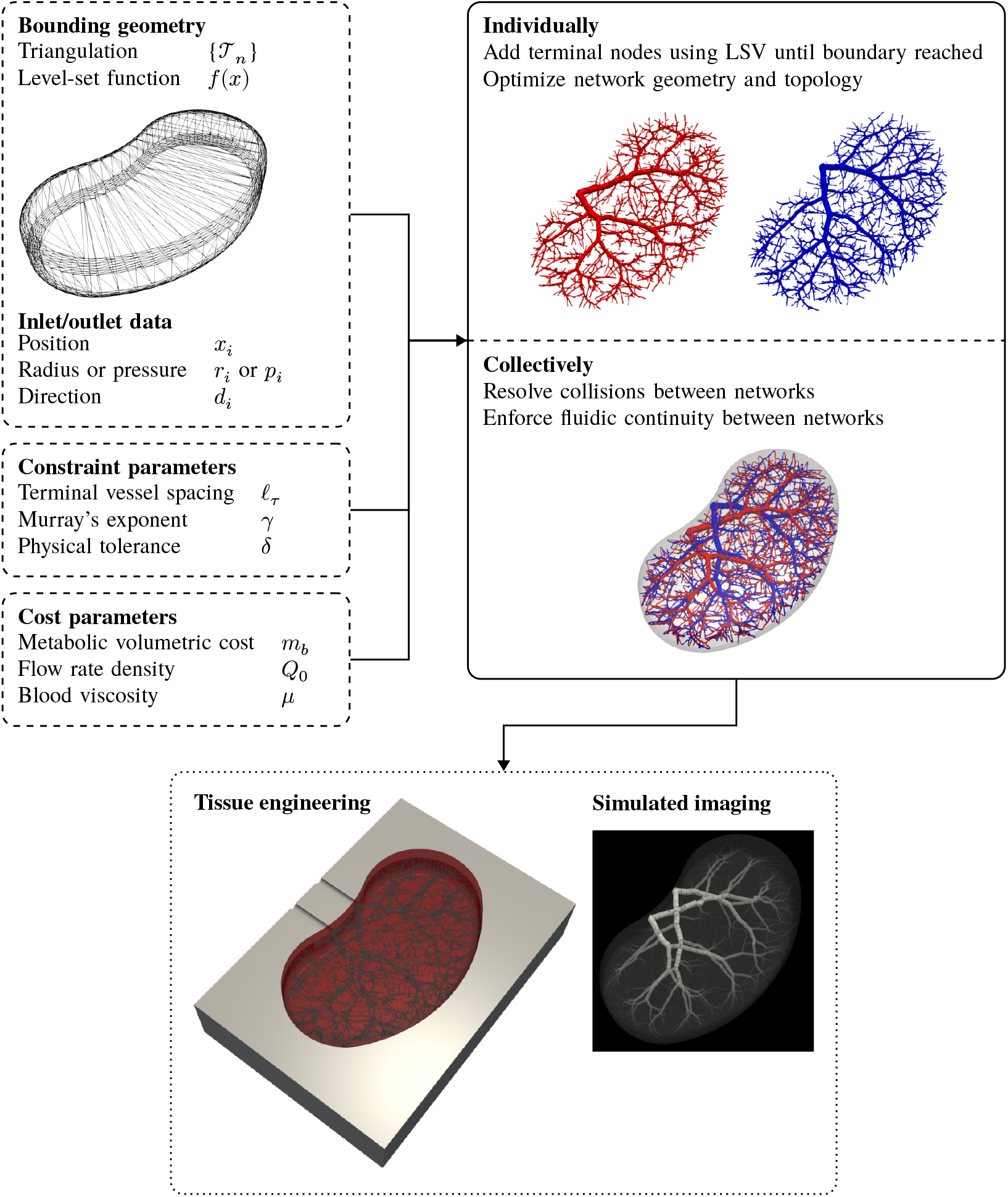
The software package *Vascular*.*Networks* contains libraries designed for the tissue engineering workflow (whose key input parameters are shown), but during validation was also integrated as one of the possible underlying growth strategies for the artificial vascular networks used as training data [38]. Consuming implementations typically iterate between individual and collective actions until convergence. In this example, the cost is dominated by the large metabolic volumetric term *m*_*b*_ (which accounts for the energy required to maintain blood volume within the vessels), so the typical early bifurcation and high asymmetry of minimal volume networks is seen.

## 2 Model Overview

### 2.1 Vasculature *In Silico*: Assumptions

#### 2.1.1 Tissue units are homogenous and self-regulating

We treat an organ as a region of space filled with nearly-identical functional structures, with a predefined length scale 𝓁 and pressure-flow relationship, which it self-regulates through the tone of the terminal arterioles. This is equivalent to the hypothesis that we could, for example, exchange two lobules of the liver or alveoli of the lungs, and they would only have to make minimal adjustments to their vessel radii in order to maintain function. We further enforce here the traditional assumption of uniform terminal pressures [12] (i.e., no large-scale interstitial flows within the organ).

For rapid assembly of tissue engineering constructs we require that the microscale self-assembles; we further expect functional, living tissue to be constantly remodelling at the microscale and to compensate for the volume lost to the major vasculature. In this framework we therefore cut off our design problem at the terminal arterioles, and any requirement for highly detailed location-specific design of the initial microscale vessels is left to be solved as a collection of separate local problems.

#### 2.1.2 Vasculature is a network of pipes undergoing Poiseuille flow

This is a well established model of flow in vascular networks (see e.g. [21]) in which a 1D approximation is used, and if nodes *i, j* are linked by a branch then the pressure-flow relationship holds:

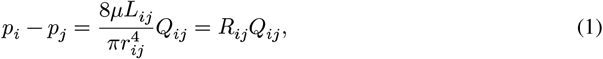

where *p* is the pressure at a node, *μ* the fluid dynamic viscosity, *L* the branch length, *r* the branch radius, *Q*_*ij*_ the directed flow from *i* to *j* and *R* the branch resistance. The mass continuity equation holds for nodes {*j*} connected to *i*:

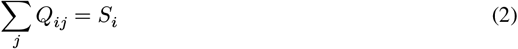

for source term *S*_*i*_ = 0 at internal nodes and non-zero at source/terminal nodes.

The pipe flow approximation is well suited to the non-pulsatile, low Reynolds number (Re) laminar flow used in tissue engineering, as the hydrodynamic entrance length for laminar pipe flow is typically stated as *L* ≈ 0.06 Re *D* for pipe diameter *D*, saturating [39, 40] at *L* = 0.06*D* below Re = 10, thus permitting vessels of low slenderness to arise from the design procedure without overly sacrificing the quality of the approximation.

### 2.2 The major vasculature is a tree

By modelling the vasculature as a tree, we either prescribe terminal flow rates *Q*_τ_ or the root flow rate *Q*_*ε*_ (in which case the *Q*_τ_ are a relative flow rate density to each other) and then at every bifurcation we set:

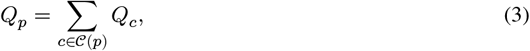

where *𝒞* (*p*) denotes the children *c* is the domain to be vascularized of the parent branch *p*. As we are feeding terminal nodes at uniform pressure, we can use the well-established notation of reduced resistance *R*^∗^ [12, 13]

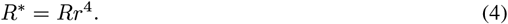

At bifurcations, reduced resistances can be propagated by considering the downstream subtrees in parallel. The total downstream resistances *R*^*d*^ therefore form a hierarchy given by

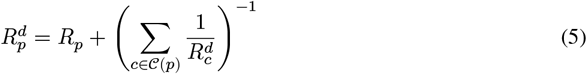

and pressure consistency between the start of sibling branches and the end of their parent requires

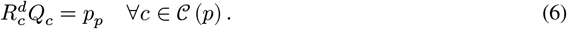

When modelling the vasculature as a tree, the child radii *r*_*c*_ can be set by their fraction *ϕ*_*c*_ of the parent radius *r*_*p*_ :

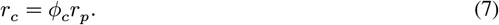

Since the terminal pressures are uniform, we note that the vector of child radii fractions 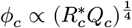 and is therefore constrained to a line. We can thus balance the tree without knowing the actual radii of the branches or the true pressure at the node, provided that the apparent viscosity of the fluid is independent of vessel radius (see 2.1.4). We then constrain this line to a point by Murray’s law [41]

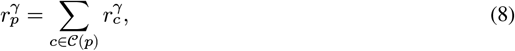

where *γ* is Murray’s constant. In terms of the vector of [*ϕ*_*c*_], this is simply a normalization in the *L*^*γ*^ norm. The value *γ* = 3 is the classical result for minimal metabolic cost, but exponents in the range 2 ≤ *γ* ≤ 3 are observed in real vasculature [22, 42].

#### 2.2.1 The working fluid has constant apparent viscosity

In pipe flow modelling, the Fåhræus-Lindqvist effect [43], which arises from the plasma/cell separation in narrow tubes [44], leads to the concept of an “apparent viscosity” that varies with tube radius *r*. This effect needs accounting for in vessels with *r* ≤ 150 µm. A radius-dependent viscosity correction prevents the efficient use of *R*^∗^ alongside the isobaric terminals assumption, as it precludes specifying flow rates solely from radius fractions and necessitates knowledge of the exact vessel radii to incorporate viscosity effects, and further requires the use of a constrained minimization approach to solve the entire problem. For instance, Schwen [29] modelled the apparent viscosity as a function of *r* using the expression:

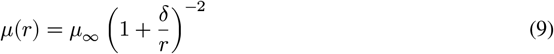

with *μ*_∞_ = 4 × 10^−3^ Pa⋅s, a common given value for blood viscosity, and cell-free layer thickness *δ* = 4.29 µm. This simplified model accounts for the decrease in viscosity in the ranges immediately below the current target manufacturing length scale, but does not account for the increase at the capillary scale as the red blood cells become increasingly large relative to the vessel diameter and the flow transitions to the plug flow regime [45].

However, in biohybrid materials used for tissue engineering, the working fluid is typically an aqueous, non-colloidal solution (i.e., cell culture medium), therefore the Fåhræus-Lindqvist effect can generally be neglected. By the time any manufactured construct would be required to be perfused with blood, vascular tone is expected to be self-regulating at this length scale, removing any requirement on our part to compensate for this effect.

Separately, an additional apparent viscosity effect arises from our use of permeable gels as the vessel wall, particularly when a cellular monolayer is not seeded onto this during manufacture. In such cases, we may deviate from the classical no-slip boundary conditions as would be seen in the real thick-walled vasculature. We note here (and derive in the Appendix A that for an idealised isolated pipe embedded in an infinite domain using the Beavers-Joseph [46] boundary condition with Saffman’s [47] approximation there is a viscosity correction of

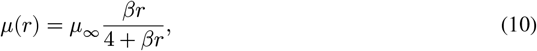

where 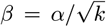, where *α* is a dimensionless material parameter characterising the slip property of the interface and *k* the bulk permeability. As the vessels we seek to design have *r* ≥ 10^−4^ m, the gels used are typically low permeability with *k* ≤ 10^−12^ m_2_ [48–51] and 0.1 ≤ *α* ≤ 10^2^ for many proposed pore geometries [52], we expect that *βr* ≫ 4 in our expected operating regime, even in the smallest vessels. Hence, we can safely use the constant viscosity model for our designs, permitting working in radius fractions alone during the design procedure.

### 2.3 The Lattice Sequence Vascularization (LSV) framework

We introduce a new algorithm designed to accommodate a wide variety of multi-scale growth processes, while ensuring uniform perfusion across arbitrary domains upon termination. The algorithm progresses through a sequence of lattices, aiming to incorporate vessels into the network by invading currently unoccupied lattice sites from their occupied neighbours, emulating the spread of real vasculature whilst the lattice sequence models a uniformly expanding growth of the tissue volume (i.e., there are no distortions arising from marginal growth at the edges as in SC). This enforces a hierarchical structure whilst ensuring that all parts of the domain are reached.

#### 2.3.1 Motivation

Ensuring that all cells are within a specified distance of a terminal channel is guaranteed by using lattices, where the maximum cell-channel distance depends on the Voronoi cell of the lattice. Whilst we have previously [30] used lattices as the terminal pattern for CCO-based strategies, real tissue does not exhibit the characteristic high asymmetry of naive CCO and other vessel sprouting approaches. During post-natal, physiological tissue growth and repair, new vessels sprout from the smaller, localised vessel networks (i.e., the capillary bed), rather than from major vessels, with ongoing remodelling regulating flow and metabolic demands. For our previous Accelerated CCO implementation [30], this requirement pushed performance towards the worst case Θ (*N* ^2^). Conversely, purely sprouting-based strategies such as SC do not have good support for remodelling, require vascular invasion over long distances, and lack a mechanism for ensuring uniform perfusion with terminal vessels.

This motivates a divide-and-conquer approach to vascularization, in which vessels sprout and invade empty space at a given terminal scale, remodel at that scale and finally subdivide into a new terminal scale.

#### 2.3.2 Algorithmic Overview

For a given lattice, we maintain a mapping of all terminal vessels located within the Voronoi cell of each lattice point, referred to as the *interior map*. In the case that only one terminal vessel exists for each lattice point, we refer to this as a *single* interior, otherwise it is *multiple*. We also define a connection pattern, which in the most restrictive case is shared faces between Voronoi cells, but can be extended to allow lattice sites with shared edges or vertices, or even allowing longer-range connections. Using this connection pattern, we visit all neighbors of the interior map, and all neighboring sites not present in the interior map become part of the *exterior map*, which represents candidate sites for growth. A pseudocode representation of the method is provided in Algorithm 1 (Appendix B).

To balance efficiency and optimality, particularly when initiating growth from a partially refined network (e.g. from scan data or L-systems), users can specify *interior modes* and corresponding *interior filter* actions (Algorithm 2). These are triggered when a multiple interior is generated. The default mode assumes that the network starts from a single vessel and has a bottleneck-free geometry, maintaining one terminal per lattice site until coarsening occurs. Alternative modes include always-single, where a selector (e.g. nearest terminal) chooses one terminal per lattice site, and always-multiple, where multiple terminals are filtered to preserve total perfusion and reduce candidate bifurcation vessels (such as by choosing extremal terminals in the directions of neighbouring lattice sites), and avoid distortion of hierarchical costs.

The main loop is performed by repeatedly calling the *Iterate* method defined in Algorithm 3. The return value of this function signals whether more iterations are possible (i.e., the exterior is not empty or there exists a more fine lattice to move to). This can either be iterated manually or handled by the *Complete* method, which enables an entire program to be loaded as a sequence of lattices and associated callbacks and executed in a single call. We then iterate three actions:

- *Spreading* (Algorithm 4): Each exterior point is visited in random order. The most optimal candidate interior point from the reachable lattice sites is selected for sprouting, and a new terminal vessel is created from it to the exterior point. After each iteration, the exterior is propagated: any point that transitions from exterior to interior is used to generate a new set of exterior points. The procedure iterates until the propagated exterior is empty, indicating that no further or successful growth is possible. To avoid overly complex higher-order splits, the default setting allows only one bifurcation per existing vessel during each spreading iteration, with the otherwise valid exterior sites which have no remaining un-bifurcated candidate vessels being reintroduced to the exterior afterwards.
- *Refinement* (Algorithm 5): Optional steps, such as topological optimization and domain boundary collision tests are executed before the algorithm progresses to the next lattice (with smaller Voronoi cells), reinitializing the interior and exterior maps.
- *Coarsening* (Algorithm 6): In domains with bottlenecks, the algorithm reverts to the previous lattice after a set number of generations. This is rarely needed in tissue engineering, as most organs are bottleneck-free. In complex cases, it may be preferable to divide the domain into sub-components and vascularize each separately.

The implementation allows user-defined actions to be executed before/after each action, and at many other points within the algorithm.

#### 2.3.3 Topological optimization via regrouping (LSV-R)

One immediate issue with this approach is that typical refinement patterns enforce similar tree structures, which are highly unnatural. Defining the normalized depth of a terminal node as in [30] by reference to the perfectly balanced binary tree (which has every node at depth log_2_ *N*), we expect that an unmodified approach for a refinement factor of 2 would lead to all terminals clustering at very similar depths over a range of terminal counts, concentrating towards the perfectly balanced binary tree as *N* increases. These structures can lead to pinning at highly complex pseudo-*n*-furcations and a corresponding loss of optimality.

To account for this, regrouping optimization (similar to that of [18]) was employed, with a modification to treat vessels as being short when they are less than 10% of their expected length according to the theory of vascular scaling (i.e., 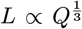) [53]. This approach is denoted *LSV-R*. Note that in order to keep the normalized depth distributions comparable for this work, we implement this regrouping using bifurcations only rather than using genuine merging/splitting to higher-degree nodes.

#### 2.3.4 Performance and scaling

By adopting the divide-and-conquer approach, LSV used with a constant refinement factor (and LSV-R with Θ (*n*) time regrouping after each lattice) is expected to achieve Θ (*N* log *N*) time in construction, with the performance depending largely on behavior near the boundary. As shown in Appendix C, this theoretical optimal performance is achieved in many practical scenarios, although we also highlight a simple case that can lead to pathological scaling. For detailed optimization at each growth stage, particularly in LSV-R, the Θ (*N*) term is expected to dominate the overall computational cost.

### 2.4 Optimization

We have previously [30] highlighted the benefits of optimizing the entire network at once rather than incrementally optimizing bifurcation positions, which is supported by recent work [31]. We report here a number of approaches we have investigated for optimizing vascular networks and expanding the number of scenarios covered by our software.

#### 2.4.1 Costs

There are a number of possible relevant costs for tissue engineering. Traditionally, just the vasculature volume, *V*, is to be minimized, as this yields the maximum amount of parenchymal tissue [12, 25, 31, 35]. However, this is usually used when operating at prescribed source-terminal pressure drop and terminal flow rates (e.g. when recreating an organ with known inlet/outlet pressures), thus the mechanical work of pumping blood through the organ, *W* (= *Q*_*ε*_Δ*p*), is fixed. In fixed root radius modes, which are more relevant when ensuring connections to pump systems, the naive minimal volume approach rewards very early bifurcations and longer, thinner vessels, which increases the pump work and root pressure required to run the device. We may consider minimizing the metabolic expenditure of the vasculature *C* (i.e. the cheapest construct to maintain), as considered by Murray [41]:

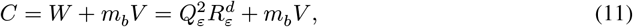

where *m*_*b*_ represents the metabolic cost of maintenance per unit volume additional blood required to perfuse the organ. Such a cost represents the additional metabolic cost of adding an organ in parallel into an existing vascular circuit, and has been considered in more recent approaches to vascular design [20].

Alternatively, we may consider maximizing the efficiency of the device *η*, defined as

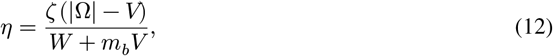

where *ζ* is the volumetric reaction rate and Ω is the domain to be vascularized. The numerator represents the reaction rate contributed by the useful tissue volume left over when the network is subtracted from a domain Ω, and the denominator quantifies the energy expenditure. Thus the ratio of the two preceding costs, *η*, captures the objective of maximizing “output” per unit energy expenditure. For open-circuit (*ex vivo*) applications such as filtration, *m*_*b*_ would be set to zero. In this equation, *ζ* may be a function of the terminal vessel spacing and flow rate, as the behaviour of the tissue depends on the microstructure. However, once a microstructure has been established, this term does not influence the optimal tree structure supplying said microscale and may be excluded from the target function. An additional correction factor may be used reflecting the lost volume that no longer needs perfusing, by setting

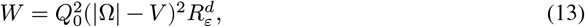

where *Q*_0_ is the volumetric flow rate density required by the useful tissue. At low terminal densities, accurate reconstruction of the downstream resistance 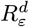 to accompany this scaling would require spatially scaling the terminal flow rates based on the local vessel geometry, as the lost tissue is not uniformly distributed (particularly when considering the major vessels). At higher terminal densities, the relative error is reduced as each terminal site feeds a smaller volume. In the case where the denominator dominates, as in real tissue (*V* small relative to |Ω| and *m*_*b*_ large [20]), all of the costs are similar and we return to the balance considered by Murray.

In previous work [30], we have shown how properties related to polynomial type costs (such as vessel length, area, volume and “hyper-volume” as considered in [54]) can be cached in the radius fraction-based framework as “effective lengths”. Recent work has used total vessel area [55] as the optimality criterion for retinal vasculature, justifying the support for generic implementations of these costs in our software. We show in Appendix D how complex costs arising from these “hierarchical cost” base components (volume, resistance, area etc.) are handled efficiently in the gradient descent computation.

#### 2.4.2 Simulated Annealing

There has been interest in the application of Simulated Annealing (SA), a well-known optimization strategy [56], to vascular design, either as the entire procedure [20, 32, 33] or as a dedicated topological optimization step in between CCO [31]. We employ a similar procedure to [31], restricting ourselves to a single early optimization pass before further growth, and using geometric optimization by gradient descent instead of making random node movements. To reduce the search space, we do not use arbitrary topological changes — we assume that the networks are nearly optimal, and instead use “promotions”, in which a node’s sibling is moved to be the sibling of its parent, thus raising the node by one in the hierarchy.

#### 2.4.3 Heuristic-guided topological optimization

As previously mentioned, we have implemented a bifurcation-only regrouping heuristic similar to the approach described by Georg et al. [18], where bifurcations are considered merged when their separation falls below a certain length. The endpoints of such an *n*-furcation are then regrouped based on their pairwise distances, and the network is geometrically rearranged to allow the nodes to separate. Additionally, we investigate a *rebalancing* heuristic, also based on node promotion, to correct highly asymmetric bifurcations at creation time [57], before resorting to more costly operations such as culling and rebuilding [30]. The heuristic follows a two-step check: First, it checks the flow rates of the child branches. If the bifurcation flow ratio *β*_*Q*_ exceeds a given threshold, we query the radii. If it is matched by a high bifurcation radius ratio *β*_*r*_, we attempt to fix this by promoting the child of the high-flow side with the highest flow rate. If the radii are biased in the opposite direction to flow rate, suggesting a long, snaking path on the low-flow side, the higher-radius side is removed and rebuilt by default.

For the software package, we have implemented a full merging/splitting procedure for *n*-furcations based on [18], where splitting is similarly based on creating an imaginary intermediate branch ι and computing a “rupture force” based on the expected cost change if this branch lengthened:

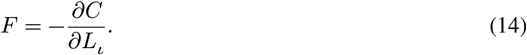

Unlike the implementation of [18], where 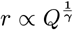and consequently the terminal pressures are not uniform, this is a more involved computation in our framework. Further, as we do not expect to create extremely high-order splits, we evaluate all possible splits for low *n*-furcations and fall back to their greedy search method otherwise. As this approach is already validated [30], we do not investigate further in this work.

### 2.5 Constraint Enforcement

Our framework resolves inter- and intra-network collisions while accounting for mesh boundaries. Boundaries are represented either as triangulated meshes (a common scenario for CAD export and medical image segmentations) or as combinations of differentiable functions — this enables the use of penalty methods in optimization rather than immediately removing branches which breach the discretized boundary. Hybrid approaches, such as fitting a set of radial basis functions to create an approximating isosurface of a mesh (e.g., by analogy to potential theory [58], which has previously been applied to myocardial shells [35]) are also possible, allowing the major vessels to be pushed back into a valid region. Typical usage employs triangulated boundaries which enable exact surface testing (although the boundary itself may be an approximation), and the optimizer can be configured with a step size controller that ensures that vessels do not form intersections with the boundary.

## 3 Results and Discussion

### 3.1 The hierarchical optimization framework enables a wide variety of costs to be considered

Figure 2 shows networks generated using an identical growth sequence, as detailed in the figure caption, with each network subsequently optimized for a different cost function. Terminal nodes are spaced 0.5 mm apart, and the root vessel has a radius 0.25 mm, giving a crown-to-stem diameter ratio of 16:1, consistent with reported physiological ranges [22]. For the cost functions, the following parameters are used: blood viscosity *μ* = 3.6×10^−3^ Pa⋅s [20]; a required tissue volumetric flow rate density of *Q*_0_ = 1.67×10^−2^ s^−1^, based on liver flow rates [59], corresponding to approximately 1 cm^3^⋅min^−1^ blood flow per cm^3^ organ volume; a metabolic cost of *m*_*b*_ = 641.3 Wm^−3^ [20] for Murray’s law type costs (*m*_*b*_ = 0 for filtration); and a domain volume of |Ω| = 256 mm^3^.

**Figure 2:**
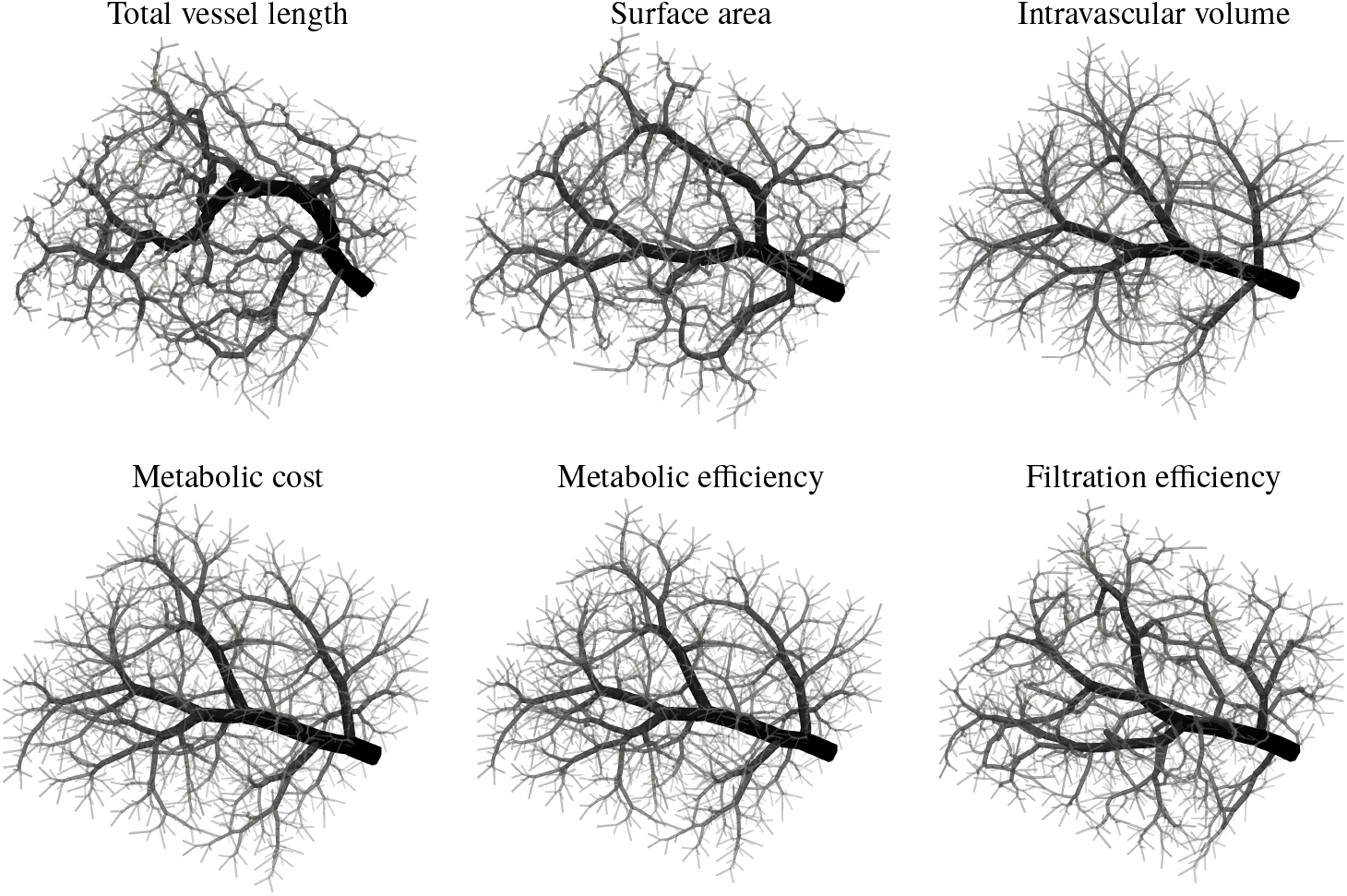
Networks generated using the same growth sequence but optimized for a variety of costs. Each network is grown from an identical terminal seque1nce: a random permutation of a 16×16×8 cuboid (*N* = 2048), with optimization at applied at every *n* = 2^*k*^ terminals for *k* ≥ 2. Terminal nodes are spaced at 0.5 mm, and the root vessel has a radius of 0.25 mm. Parameters used in the cost functions: *Q*_0_ = 1.67 × 10^−2^ s^−1^, *μ* = 3.6 × 10^−3^ Pa⋅s, *m*_*b*_ = 641.3 W/m^3^ (metabolic), *m*_*b*_ = 0 (filtration), and |Ω| = 256 mm^3^.

It can be seen that we reproduce visually similar patterns to those reported by [54] for simple polynomial costs (length, area and volume), and to those in [31] for metabolic cost. Under physiologically realistic parameters, Figure 2 demonstrates that networks optimized for metabolic cost *C* and metabolic efficiency *η* differ only marginally. This is expected when the intravascular volume *V* is small relative to the domain |Ω|, implying that the blood maintenance cost dominates. A visual comparison of the networks optimized for minimal intravascular volume, minimal metabolic cost and maximum metabolic efficiency is shown in the Appendix Figure D.1.

A full comparison of our simulated annealing approach to that of [20] is presented in Appendix E Figures E.1, E.2 and E.3, showing superior results in terms of metabolic costs for an approach similar to [31] on the standard test case of [20]. Since the root vessel structure dominates the overall network cost [14, 54, 57], while the influence of “deep” nodes is relatively minor [30], constructing the full network before applying simulated annealing, as in [20], is inefficient. Instead, simulated annealing should be applied early, when each terminal vessel represents a volume that will eventually contain many other vessels. As demonstrated in the example presented in Appendix Figure E.3, this approach converges to similar structures using significantly fewer iterations (10^6^). Nevertheless, simulated annealing remains an extremely computationally expensive approach. Designs within 1% of the simulated annealing output can be achieved at far lower computational cost by generating thousands of candidates using biologically-inspired growth methods (at this scale, the low overhead of ACCO makes it extremely fast) and selecting the best, a process that is trivial to parallelize. As such, simulated annealing should only be used for cases where the computational investment is justified, such as designs intended for mass production, while heuristic-based optimization is more suitable for prototyping.

### 3.2 LSV scales nearly linearly and regrouping optimization results in self-similarity

Fig. 3(a) shows build time *t* and fitted scaling laws *t* ∝ *n*^*α*^ and *t* ∝ *n*^*α*^ log *n* for the basic LSV actions (spread/refine) i.e., without any post-spread optimisations to the network. Results are shown for both the default setting (referred to as “multiple build”) and an optional configuration that restricts each spreading iteration to a single bifurcation from each existing terminal (“single build”). This comparison highlights that, in the case of a cubic lattice, the single build incurs a performance penalty due to requiring three successive doubling steps rather than completing the full eightfold refinement in a single iteration. Additionally, we also note the relative variance at low values of *n*, which is likely unavoidable when running on consumer operating systems due to the short time taken relative to process scheduling. Also shown is an adaptation of ACCO, with a “staged growth” approach based on LSV, where for each lattice all newly added terminals are shuffled and then appended to the queue. It can be seen that LSV becomes significantly more efficient at larger scales, with the crossover point for the slower single build configuration at around *N* = 300. In particular, the near-linear scaling allows the growth of highly-detailed vascular networks in the order of minutes, spending far less time on the finer vessels (which impact the overall cost of the network less) than ACCO, whilst the algorithmic overhead at smaller scales and need for regrouping suggests that ACCO should be used in the earlier stages, where highly asymmetric branching is unlikely to occur.

**Figure 3:**
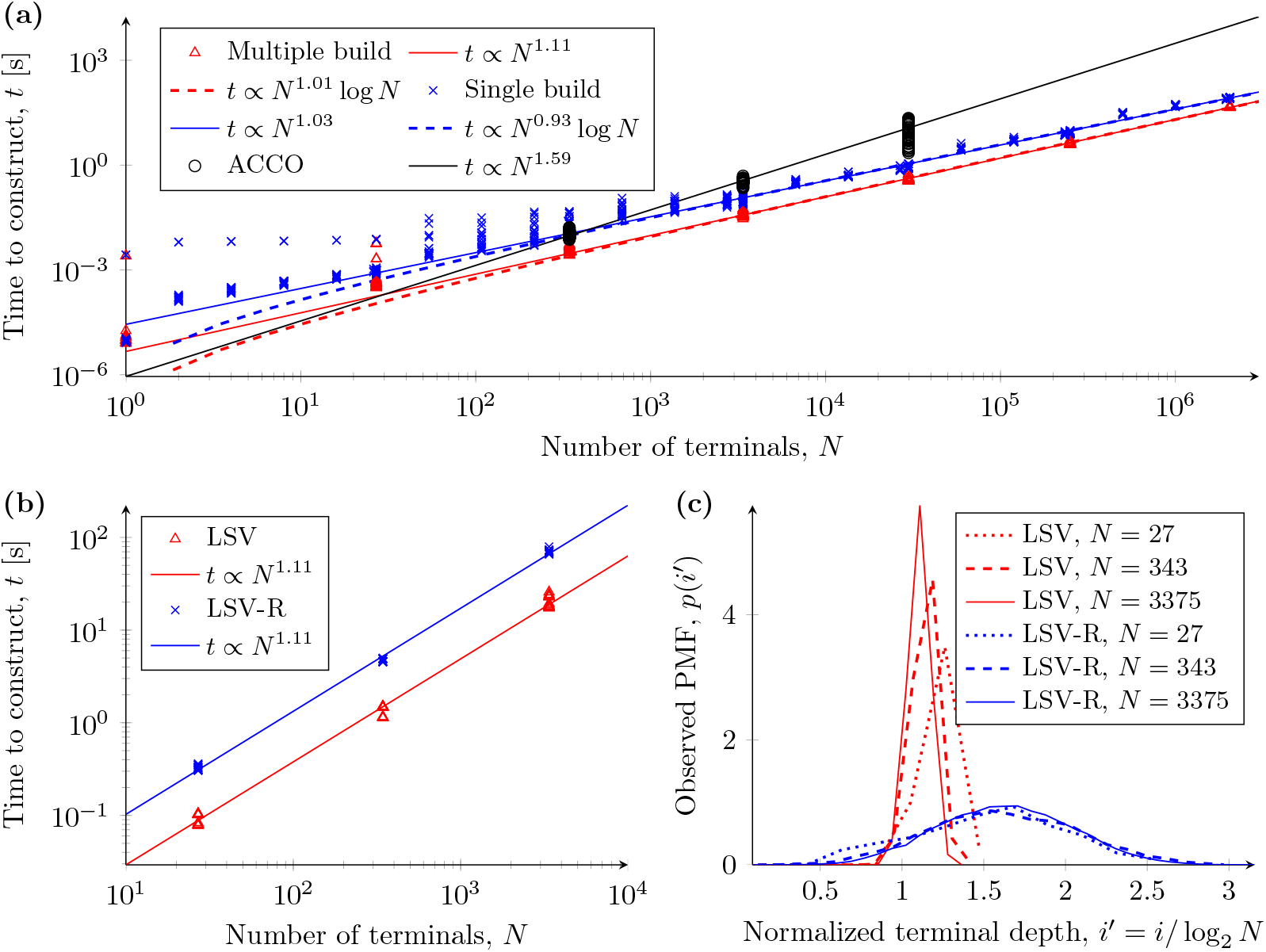
Build time *t* and terminal depth distributions *p*(*i*^′^) for LSV and LSV-R. Data taken from 25 instances of each growth method. (a) Time taken t1o build a network using LSV as a function of final terminal count *N*, and an estimate of power law scaling. Also shown are timings of staged ACCO using the same lattice sequence. (b) Time taken to build a network using LSV-R (LSV with post-spread optimization) as a function of *n*, and an estimate of power law scaling. Note that there is a high variance in times taken, as the number of iterations required for completion at each level is variable and each iteration has high computational cost. Data taken from 25 instances of each growth method. (c) Terminal depth probability mass function (PMF) *p*(*i*^′^) using LSV and LSV-R, where *p*(*i*^′^) represents the fraction of terminals at normalized depth *i*^′^.

Fig. 3(b) shows that when optimization is incorporated after each iteration, the cost of this step becomes the dominant factor at the sizes currently of interest. This is even more pronounced for LSV-R, whose cost is primarily determined by the number of iterations of regrouping performed at each length scale. In practice, when both stabilising behaviours are activated (only one bifurcation from each branch per iteration and regrouping optimization), we should allocate linear time for growth of vascular networks at typical tissue engineering scales, and treat the LSV time itself as negligible.

In order to compare terminal depth distributions, using *p*(*i*) to represent the fraction of terminals at depth *i*, the same normalization as in [30] is used. Since the perfectly balanced tree will have all terminals at depth log_2_ (*N*), the comparable distributions are referred to as *p*(*i*^′^), where *i*^′^ = *i*/ log_2_ (*N*).

In Fig. 3(c), the LSV approach, for a refinement factor of 2, shows concentration toward the theoretical perfectly balanced binary tree with increasing *N* (a pathological state), whereas LSV with regrouping (LSV-R), enables the resolution of higher order splits. Importantly, the terminal depth distribution is spread over a broader range, and this distribution remains unchanged over several orders of magnitude of *N*, suggesting a degree of self-similarity across length scales, concordant with some of the theoretical scaling considerations of vasculature [22, 53] and also Murray’s law.

LSV, when combined with optimization methods that are able to make modifications to the local branch topology, is therefore both a suitable candidate for early vessel growth as well as for ensuring complete perfusion in the later stages of vascular design.

### 3.3 LSV enables functional microscale structures to be supplied without risk of short circuit

For tissue engineering, a key challenge is replicating tissue functionality at the microscale. While this will typically need to be performed by self-organization due to manufacturing limitations (and the tendency of tissue to reorganize itself regardless), there are instances where it may be beneficial to provide specialised structures for the initial state. The simplest example of this is the insertion of a lattice of vessels at the smallest manufacturable radius. Figure 4(a)(i) shows a 3D-printable meso-capillary lattice that could support a typical biohybrid material such as a densely populated cell-laden gel. In this example, capillary lattice edges which intersect major vessels or form dead-ends are removed (Figure 4(a)(ii)) and connections are only permitted to terminal arterioles/venules (Figure 4(a)(iii)). This enables a biomimetic spacing and radius in the meso-vasculature whilst still ensuring correct fluid transfer between arterial and venous trees, as well as providing a template for capillary bed formation and reducing the distance required for cells to invade from channel walls into the bulk. Further, such structures may eliminate the need for support waxes (as in the case of the major vessels alone) to be used in the printing process, thus eliminating the risk of template breakage during de-waxing. In our framework, the development of the major vessels and the dense capillary bed are separated, allowing the capillary bed to be “streamed” in chunks for manufacture, saving computational resources.

**Figure 4:**
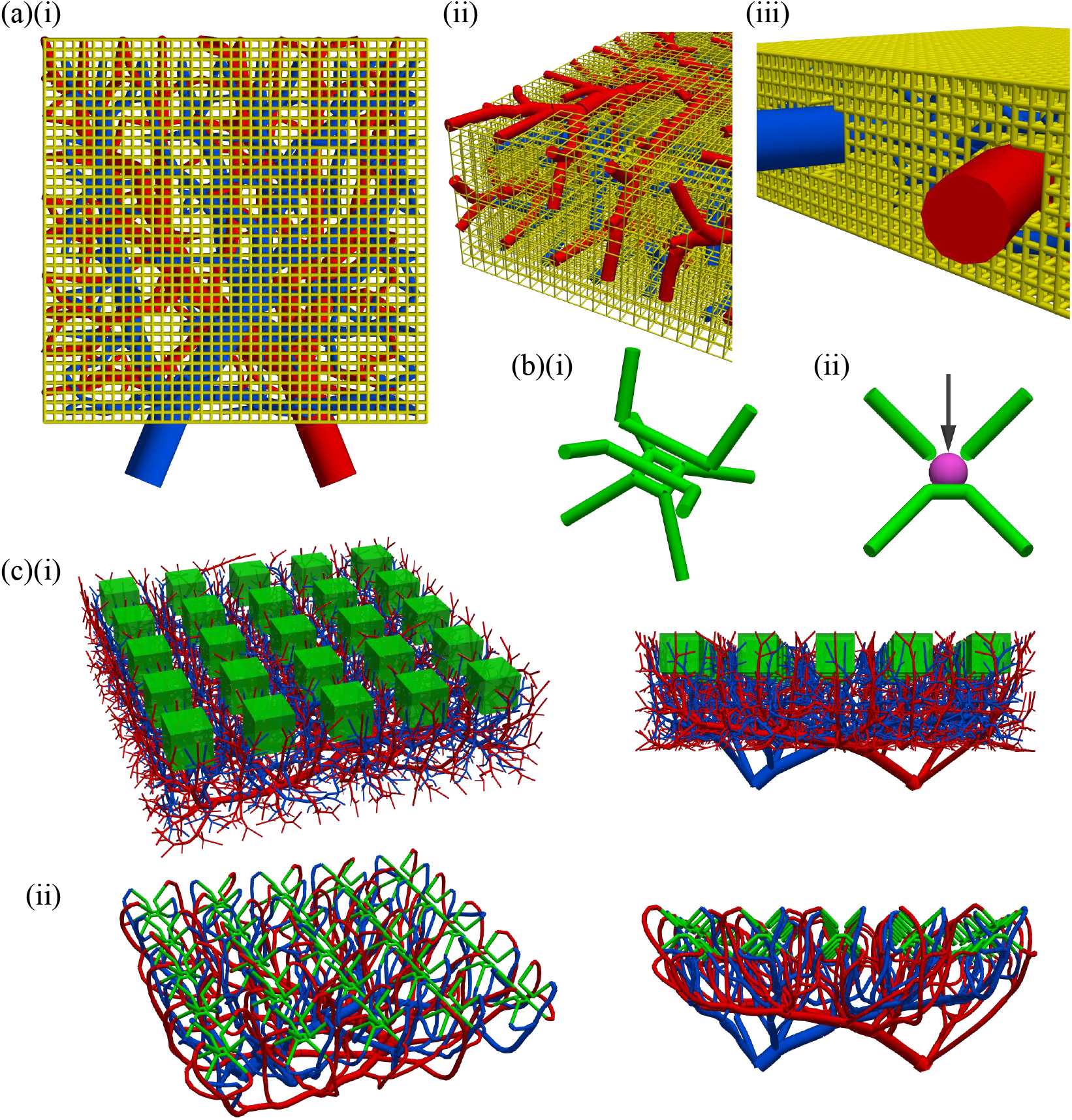
Examples of functional units supported by the software package: a capillary lattice and an organoid trap. (a)(i) A 3D-printable meso - capillary lattice, with channels of diameter 200 μm spaced at 500 μm. Terminals are spaced at 2 mm with arteriovenous offset of 1 mm, leading to a visibly over-vascularized construct. The connection policy permits intersections with segments near the end of the terminal vessels, (ii) permitting some connections to terminal arterioles/venules, within one capillary lattice spacing of the intended terminal and (iii) clearing space around the major vessels. (b) (i) An organoid trap is shown in isolation, with the intended loading direction and containment of an organoid in (ii). (c) The networks were grown using LSV with the insertion locations kept clear (i), with a terminal density far greater than strictly required to ensure existence of a valid connection; the majority of these vessels were trimmed after the functional units were inserted as shown in (ii).

The feasibility of manufacturing complex discrete functional structures with multiple fluidic paths, essential for the proper functioning of organs such as the lung, kidney, and glandular tissues, and equally relevant to the design of parallelised microfluidic mixers and reactors, is demonstrated in [60]. It is therefore of interest to provide software functionality associated with tiling these functional units throughout a given tissue volume.

In this work, we consider the example of a simple “organoid trap” with 4 inlets and 4 outlets, designed to contain a cell-dense organoid at the limit of oxygen diffusion (typically around 500 µm diameter) and metabolite diffusion (typically around 150 µm) [61, 62]. The organoid trap has direct channel connections, a supporting platform and bendable top rails to allow insertion and containment of the organoid (Figure 4(b)). For visual simplicity, we do not develop 4 distinct fluidic network pairs; instead, we designate alternate corners as being arterial/venous. (There would be no algorithmic adaptation required to generate distinct networks for different fluidic tracks, which could be used to establish solute gradients across the organoid if desired.) After tiling the desired functional structure and creating a boundary mesh representing the exclusion zones, we use LSV with a terminal resolution fine enough to ensure that all gaps are filled (Figure 4(c)(i)), before selecting the optimal candidate branches and making connecting vessels to each inlet/outlet and trimming the remaining vessels, leaving a correctly supplied array of functional units (Figure 4(c)(ii)).

### 3.4 Additional enhancements improve biomimicry and enforce physiological constraints

Figure 5 shows a number of biomimetic enhancements that have been integrated into our software. Firstly, we adapt the classical approach to radius splitting between parent and children. Murray [41] reasoned that a network optimizing the total energy costs required both to pump fluid through the vasculature and to maintain the volume of blood would yield a splitting rule (8), with *γ* = 3. However, in real vasculature it is seen that 2 ≤ *γ* ≤ 3, with lower values in the larger vessels [22]. Therefore, committing to a single exponent for a large network will increase biomimicry in one region at the expense of biomimicry in another. Additionally, this can lead to issues with manufacturability, where radii decrease too rapidly at low *γ* (as the slowest decay occurs with symmetric splits, where *ϕ*_*c*_ = 2^−1/*γ*^) and fall below the minimum feature size; conversely, at higher *γ*, root vessels do not decay rapidly enough and can lead to excessive tortuosity following collision resolution.

**Figure 5:**
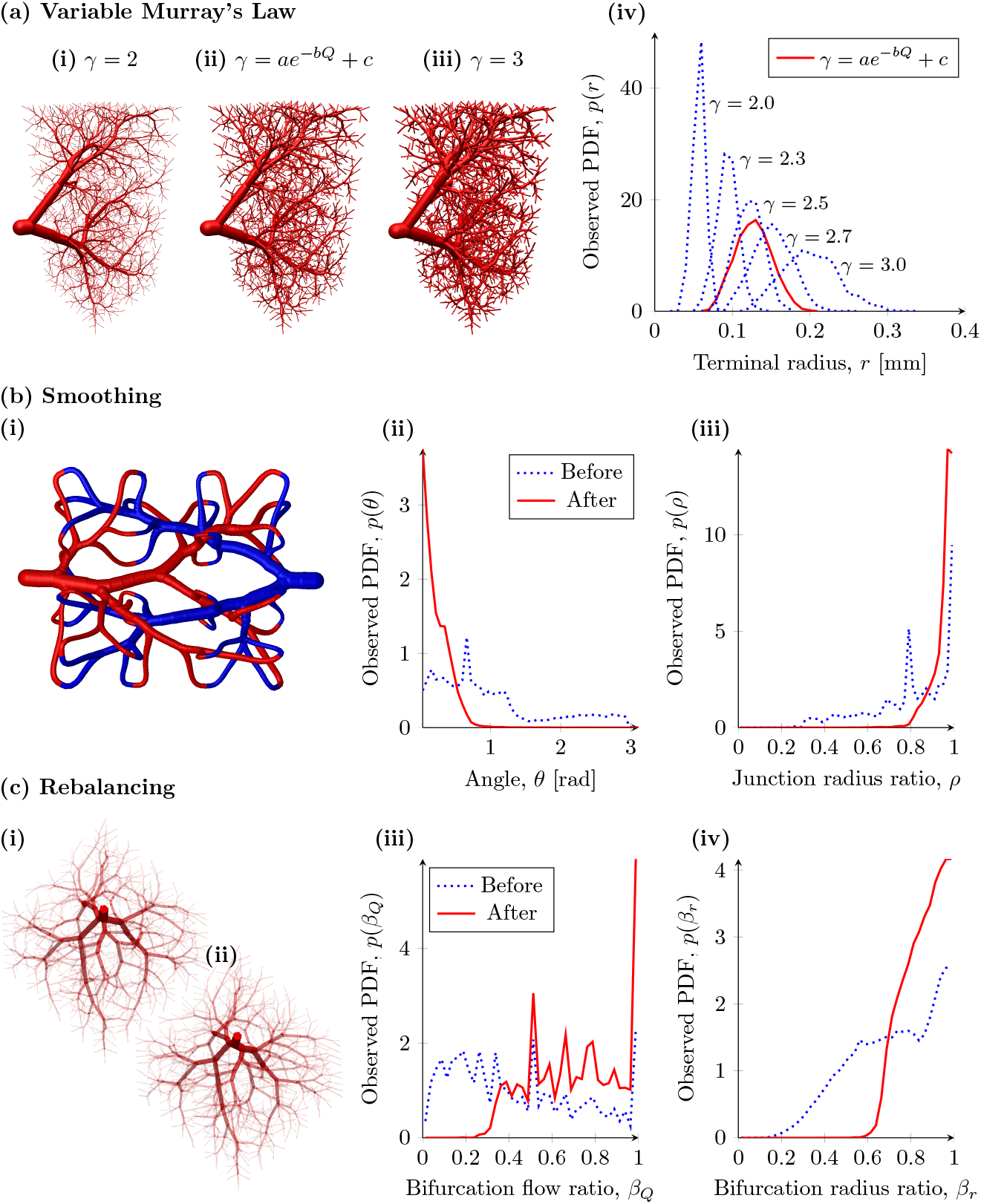
The incorporated features for increasing biomimicry and enforcing empirical physiological constraints. **(a)** Variable Murray’s law. The same1network with *N* = 2000, *Q* _*term*_ = 1 mm^3^s^−1^, *r*_0_ = 3 mm and terminals spaced at 5 mm, with different splitting laws applied. In (ii), we set *a* = 1.05, *b* = 2.82 × 10^−3^, *c* = 2.00, which was achieved by fitting to the extrema: *γ*(*Q*_τ_) = 3 (terminal), *γ*(*Q*_*ε*_) = 2 (root), and an (arbitrary, for demonstration purposes) intermediate point, 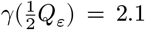. (iv) Changing γ as a function of *Q* allows improved biomimicry at all scales whilst still meeting design specifications. **(b)** (i) Vessel smoothing at junctions. The impact of smoothing on (ii) junction angles and (iii) radii ratios is illustrated using probability density function (PDF) plots. Note the elimination of sharp angles and radius transitions. **(c)** Bifurcation rebalancing. In an extreme example, we set the critical ratios 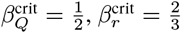. An example of its action can be seen in (i)–(ii), as it resolves a 4-way split at the root into two levels of balanced bifurcations. (iii) The most extreme flow ratios have been adjusted upwards and (iv) all radii ratios are above the critical value as shown in the PDF plots.

To address this, we permit Murray’s exponent *γ* to vary. Using flow rate *Q* as a proxy for position in the tree, we define

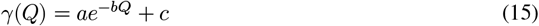

and evaluate *γ* at a branch node. In this formulation, *γ* increases as *Q* decreases. For example, when applied to the model in Figure 5(a), the variable exponent *γ* model (Figure 5(a)(ii)) is seen to have fast inlet decay at the radius due to the lower *γ* (which is also desirable for manufacturing the maximum amount of useful tissue), whilst not decaying excessively at the terminals. The function parameters (*a, b, c*) were set by fitting to the extrema and an intermediate point, thereby enabling control over radius decay in the intermediate vessels to meet specifications at the inlet and terminals. To achieve a similar distribution of terminal radii for the same inlet radius with using the fixed exponent model would require *γ* ≈ 2.5 (Figure 5(a)(iv)). Note the similar root vessel structure between (i) and (ii), and that the terminal vessels of (ii) are closer to those in (iii), showing slower terminal decay. While a constant Murray’s exponent allows higher-order splits to be simulated using only bifurcations, Appendix F describes how the same can be achieved with a variable exponent.

Secondly, we enable the smoothing of networks to eliminate sharp corners and abrupt radius transitions at branch nodes which helps to improve the quality of the pipe flow approximation to real flow rates, and when used to generate training data for medical imaging, makes the vasculature look more natural. Angular smoothing is achieved by modifying the gradient descent minimizer to take the total energy associated with a linear and angular spring model of the vessels, whereas radius transition is performed using a Laplacian-type smoothing (see method of [63, Ch. 5.4]), with renormalization after each iteration to ensure that the resistance of the branch remains unchanged, i.e.:

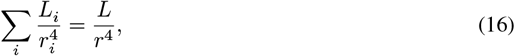

where *i* indexes the segments of a branch. We note that the angular smoothing will tend to reduce optimality for most costs by introducing additional length, and that the radius smoothing will always reduce optimality for polynomial terms in cost expressions - see Appendix G. In Figure 5(b)(i)), we demonstrate a simple tissue support design compatible with current 3D printing methods, with all vessels exceeding 200 μm in diameter. The design assumes a working fluid of water with cellular density 1/1000th that of real tissue, allowing optimization solely for maximum useful tissue since the pumping work for such low required flow rates will be negligible. A realistic terminal spacing of 4 mm is used, which for a 4×4×2 terminal layout requires an inlet diameter of ∼1 mm (suitable for connection to a standard (ISO6009) 17G hypodermic needle, which has inner diameter 0.042 in = 1.0668 mm). Two iterations of smoothing were used with a linear and angular spring constant of 1, with a boosting factor of 5 for the terminal vessel end segments to ensure separation; and 50 iterations of radius Laplacian smoothing with renormalizing were used in the final stage of collision resolution, which led to a mean volume increase of 9.1% compared to the uniform profile. This can be seen to give highly smoothed vessels with no sharp angles or sudden changes in radius. Figures 5(b)(ii)–(iii) show the concentration of junction angles around 0 and radius ratios around 1 arising from this scheme, when tested on 100 instances (the slight peaks before smoothing arise from the consistency of LSV-R when used on such a small target (*N* = 32), recreating similar structures each time).

The final modification presented here is the enforcement of bifurcation symmetry through topological rebalancing optimization (see 2.3.3). The *rebalancing optimiser* promotes or demotes nodes based on their flow and radius ratios at a bifurcation, with the option to cull if necessary. Real tissue has been noted to contain radius asymmetry ratios of 5:1 [57], with no quoted values for flow rate. In Figure 5(c), we show an extreme example of enforcing bifurcation symmetry, with target flow rate ratio 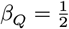 and radius ratio 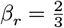, and allowing 5 iterations of geometric optimization interspersed with rebalancing iterations. Whilst there is often no solution for flow rebalancing at such an extreme critical value, we find that we are able to enforce the radius imbalance criterion very closely without needing to cull large sections of the network. Figure 5(c)(i)–(ii) illustrate the capacity of this approach to rearrange “pinned” bifurcations (e.g. near the root vessel) into a more optimal topology that the geometric optimizer can subsequently refine. Figure 5(c)(iii)–(iv) show the distribution of bifurcation ratios before and after the procedure. In this case, we can enforce a vascular design that is well within biologically observed asymmetry ranges.

### 3.5 Human-scale vasculature and supporting non-vascular networks can be efficiently generated using LSV

Many geometries cannot be efficiently represented using solid constructive geometry built from simple analytic shapes. In addition, realistic vessel networks must often be generated within patient-specific domains reconstructed from imaging data. For these reasons, we focused on supporting triangulated boundary representations. This is handled primarily through terminal-pair predicates. Supported modes include checks that: (i) the line segment connecting the candidate terminal pair or (ii) the triangle bounding the candidate bifurcation triad does not intersect the boundary. Simpler tests are also available, such as verifying that the terminal lies within the interior of the geometry, though this may require post-processing to cull terminals that intersect the boundary after spreading.

A test-case of particular interest is the liver, a complex organ with two supply networks, the hepatic artery (HA) and portal vein (PV), the hepatic vein (HV) and biliary tree (BT). Figure 6 shows an example of vasculature grown in a human liver reconstructed from scan data, which used the line segment condition for early vessels at large stride (as the triangle condition was too strict and prevented growth), before switching to the triangle condition. Extra complexity arises from the portal triad (PT), in which the HA, PV and BT have similar structure, acting as a single tree with three distinct channels; further, the desired functionality requires recreating the lobular structure of the liver at the finest scales. This, coupled with the large physical size of the liver, complex bounding geometry and clinical need for replacement tissue, makes it a demanding and popular test case for vascularization software [25, 28, 29, 33].

**Figure 6:**
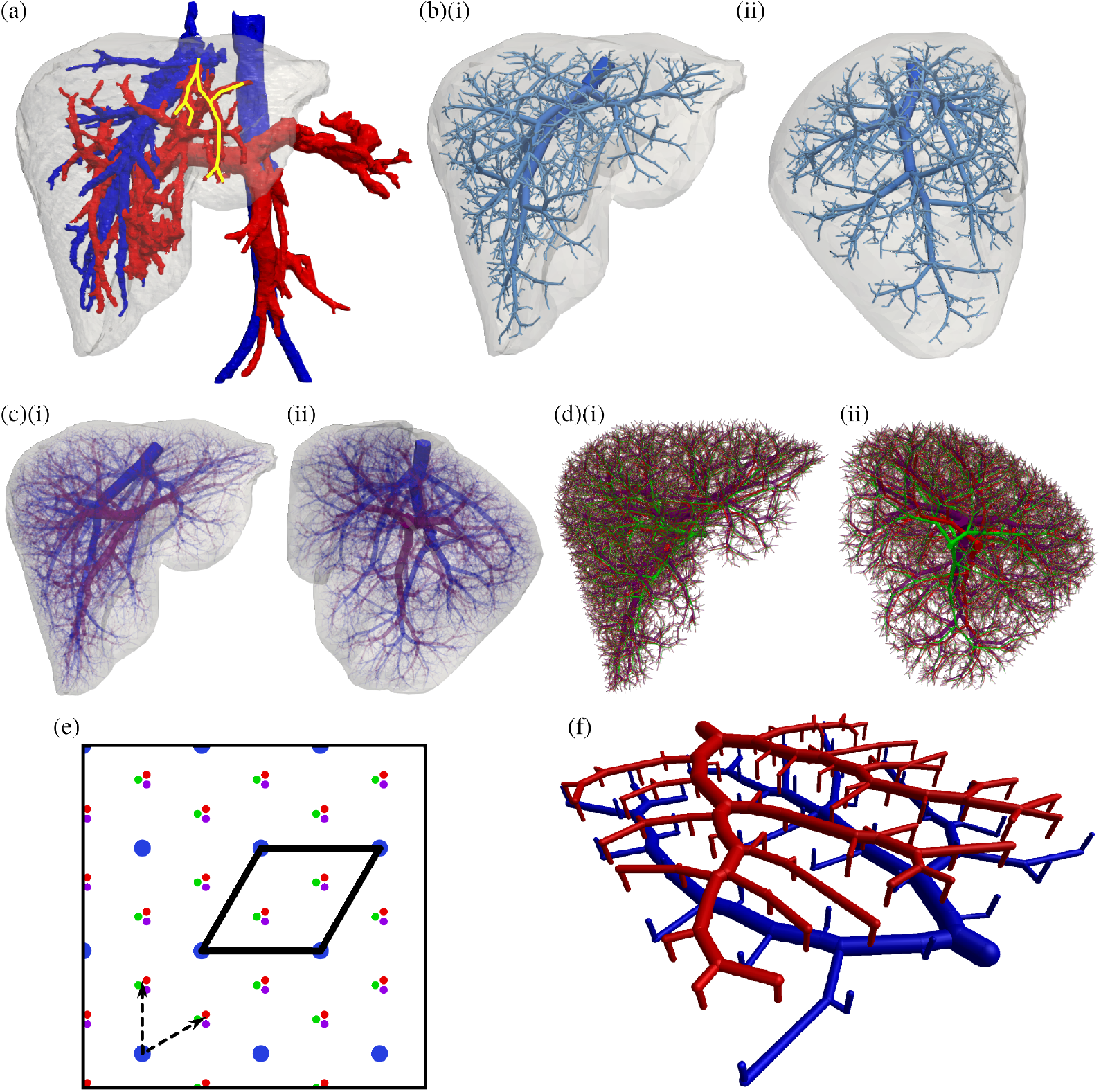
Design of metabolically-optimized vasculature in a human liver geometry, using LSV-R. (a) A human liver grown in a real bounding geometry, taken from Embodi3D, with hepatic vein (plus external section of inferior vena cava) coloured blue and portal triad (plus external section of abdominal aorta and coeliac trunk) red. Copyright 2021 Selami Ekinci, released with permission under a Creative Commons Attribution 4.0 International License. The bounding geometry and segmented vessels were used to set approximate inlet/outlet positions and radii. Note a likely venous vessel that has been misattributed to the portal vein due to a break in the segmentation (highlighted yellow). (b) A completely synthetic hepatic vein, with only vessels of Strahler order greater than 2 plotted. (c) The hepatic vein (blue) and portal triad (purple) pair rendered in front (i) and side (ii) view with the bounding mesh, with vessel opacity a function of radius for clarity. (d) The portal triad, split into its constituent networks with collisions resolved, with hepatic artery (red), portal vein (purple) and biliary tree (green) offset by 20 mm at the root, with lower offset at greater logical depth. To prevent visual clutter, vessel opacity is a function of radius. (e) The lobular tiling of the liver. The lobules are arranged in a hexagonal tiling, with the portal triad and hepatic vein columns being in a 2:1 ratio. A possible unit cell is highlighted, and two possible basis vectors are shown. (f) A liver-on-a-chip design for a single layer of hepatic lobules, with columnar structure added.

Defining the networks using a triangulated human liver bounding geometry, the source nodes and inlet radii for each network were placed by manually sampling points on the segmented vessels (Figure 6(a)). The HV and PT were grown using LSV-R with the target cost being minimal metabolic expenditure as in Section 3.1 (Figure 6(b)–(c)), after which the PT was split into its constituent networks using a radius-dependent offset (such that the root vessels were sufficiently spaced whilst the terminal vessels converged), and collisions were resolved between all networks (Figure 6(d) shows portal triad only due to vessel density). Of note is the reconstruction of similar vascular features to the real liver, such as the portal splitting horizontally upon entry and the hepatic vein splitting into three major branches, although the exact anatomy is not emergent and we would not expect a simple energy expenditure argument to completely define the liver vasculature. In particular, the “solidification” of vascular structure as the major vessels form and non-uniformities in organ growth will contribute to non-optimality in the final configuration.

The functional structure of the liver arises from its lobular tiling of PT and HV vessels, in which a hexagonal tiling emerges with a 2:1 ratio of PT to HV columns. Figure 6(e) shows the lobular structure, a possible pair of unit vectors and a unit cell of the lattice. It is clear that the HV columns can be grown using LSV on a sub-lattice of index 3, whereas the PT can be formed by knocking out terminal sites corresponding to HV sites, either using an external predicate or by trimming the vessels post-growth, enabling the development of liver-on-a-chip designs (Figure 6(f)) when confined to a single plane. The liver terminal lattice employed here is described in Appendix H.

## 4 Fabrication of Biohybrids

Complex vascular architectures enable the fabrication of biohybrid devices with higher-order, biological vascular features – not currently producible via cellular self-organisation principles - and those which have no direct biological equivalent which can be directly engineered for function and resistance to damage and disease. Complex vascular networks can be explicitly engineered for fault tolerance, providing redundant flow paths and a faster response that preserve perfusion after localised damage or occlusion, thereby avoiding ischemic injury and loss of function. Redundancy is an important feature of the native, biological vascular system: this includes arterial anastomoses (e.g. Circle of Willis), collateral circulation (e.g. collateral coronary vessels, cerebral leptomeningeal vessels, mesenteric vessels), dual supply vessels (pulmonary and bronchial arteries), and interconnected vessel plexuses (e.g. vertebral veins, pelvic veins) which are resistant to hierarchical vascular failure [64–68]. Finally, microvascular capillary beds inherently provide robustness through dense and redundant vessels, meaning local failures do not disrupt tissue functionality [69]. Using biological vessel systems as a guide, such fault tolerant vasculature can be designed into biohybrid tissue devices, greatly improving their resilience, alongside non-biological advantages such as hyper-perfused tissues or parallel delivery of metabolites and drug delivery. Finally, engineered vasculature decouples transport performance from cellular function – for example not relying upon angiogenesis for optimised oxygenation – reducing variability and providing predictable performance across devices.

Fabricated vascular networks can be designed within a range of synthetic biomaterials, imparting non-biological benefits to a living tissue and make such devices more resistant to failure. For instance, smart polymeric biomaterials can be developed which provide a self-healing nature, comparable to self-sealing concrete. Using synthetic materials instead of ECM enables the engineering of biological tissues which are resistant to mechanical damage (e.g. synthetic cartilage, skin). Synthetic biomaterials can be made intrinsically resistant to disease processes including thrombosis - through non-fouling surfaces and nanoparticle release; infection through functionalised polymers with localised antibiotic release; and tumour growth and angiogenesis using functionalised polymeric surfaces for localised drug release. Additionally, biohybrid devices with engineered vasculature can operate under non-physiological conditions, including variable temperature and chemical environments. Finally, the design of complex vascular architectures allows modular replacement or component swapping of biohybrid devices [70–72]. A reproducible vascular design means that specific regions of a damaged tissue or device can be replaced without the need for biological cellular remodelling. Current bioprinting methodologies that support vascular network fabrication typically involve photocrosslinked and smart bioinks (e.g. digital light processing, stereolithography, volumetric bioprinting) providing a synthetic nature to tissue engineered devices . Indeed, such devices have been made consisting of a biological hydrogel pocket (e.g. made of fibrin gel) surrounded by a synthetic biomatrix supporting complex vascular designs [60]. Therefore, such biohybrid devices provide early examples of biohybrid devices which may enable the functionalities described above.

Beyond the need for living tissues, there are a wide range of biohybrid applications for engineered vascular network designs [73]. This includes soft robotic functions for actuation, sensing and control; the incorporation of biological cells into such devices as biochemical sensors (chemical, innervation) and actuation (muscle tissue actuators), which require metabolite delivery and exchange; rapid, controlled and simplified vascular designs for specific devices; and high throughput bioprocessing systems which take advantage of the 3D nature of the vascular network, compared to typical 1D channel screening. Finally, engineering vasculature can function as a highly efficient heat dissipation mechanism for bioelectronic devices.

## 5 Conclusions

In this work, we present Lattice Sequence Vascularization (LSV), an iterative growth algorithm developed to design vascularized biohybrid tissue constructs. LSV enables growth through bottlenecks, supports major vessel development in non-convex geometries and ensures complete terminal perfusion with good computational scaling. The algorithm relies on a network optimizer capable of modifying both network topology and geometry to generate high-quality networks. By interspersing optimization steps regularly with growth, the required optimization effort remains bounded.

LSV operates through multiple stages: *growth*, where new vessels are added to the tree; *optimization*, which adjusts geometry and topology to minimize competing cost functions while preserving perfusion; *constraining*, to prevent premature intersections and enforce boundary limits; *biomimicry*, to enhance physiological relevance; and finally *functionalization*, where multiple trees are connected at terminal vessels through a functional structure. Major contributions include improved optimization routines; enhancements also applicable to the tissue-engineering-adapted CCO algorithm, enabling the generation of organ-scale vasculature with unprecedentedly low computational cost; and the integration of functional structures to achieve enhanced biomimicry across multiple length scales.

## 6 Acknowledgements

AAG was supported by the Engineering and Physical Sciences Research Council (EPSRC; EP/N509620/1). We also acknowledge support from the Wellcome Trust (DRP 226795/Z/22/Z).

## Appendices

### A Apparent viscosity in hydrogel constructs

Considering the first-order coupling of Beavers and Joseph [46] at the boundary:

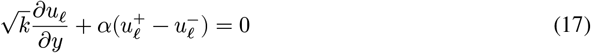

where 𝓁 is the segment lengthwise direction, *y* the wall normal direction, *α* a dimensionless material parameter characterising the slip property of the interface and *k* the bulk permeability, and the superscripts signify that this is the tangential velocity slip on either side of the interface. By neglecting the effect of other channels in the vicinity of a channel interface, we consider the continuity of pressure across the free bulk and interstitial flow, in which Darcy’s law holds:

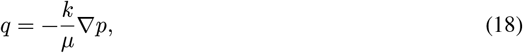

where *q* is a bulk flow rate per unit area (volumetric flux). This is an adaption of the reasoning of Beavers and Joseph in their seminal paper [46], and we note that the difference between the apparent and physical velocities is linearly related through the porosity of the medium, which may be absorbed into the material constant *α*.

It is well-established that the solution for laminar flow in an axisymmetric cylindrical channel is a parabola *u*_𝓁_(*y*) = *a* + *by*^2^. Let the interstitial flow be given by *q*, and 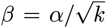, then we find at the boundary:

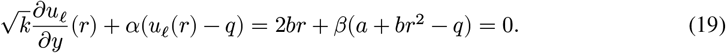

We may combine this with the relationship for *Q*(*a, b*) to yield a linear system:

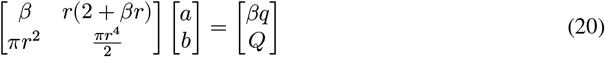

which has determinant 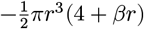, and so is always solvable for pipes with non-zero radius. At the boundary, we have the condition:

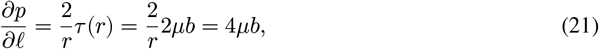

and so we solve for *a, b*, rearranging to get familiar powers of *r*:

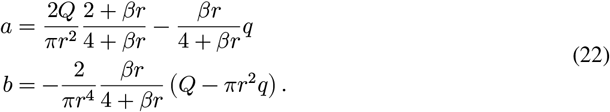

Inserting this expression into (21), and combining with Darcy’s law gives

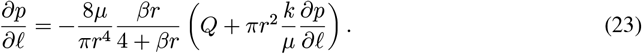

In the case where bulk flow is negligible (Saffman’s modification [74]), setting *q* = 0 in (22) yields

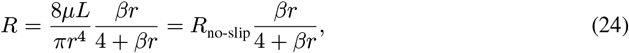

the product of the classical resistance and a factor, dependent on material properties and radius, which approaches 1 as permeability approaches 0 (classical no-slip condition) and 0 as permeability approaches infinity (implying that the behaviour is as though there is no pipe and the entire domain is conducting the fluid). We can also examine the velocity profile as a check: as *β* becomes large we recover the classical pipe profile, and as it becomes small we reach a constant profile at the mean flow. We also find that the impact of this term is felt most in the smaller branches: for increasing *r* we return a factor closer to unity.

### B LSV Algorithms

#### Algorithm B.1

The utility functions provided. Note that in practice, these methods have overloads that allow them to operate more efficiently in the single-interior case (where the interior map *I* always returns a single-element set or ∅).

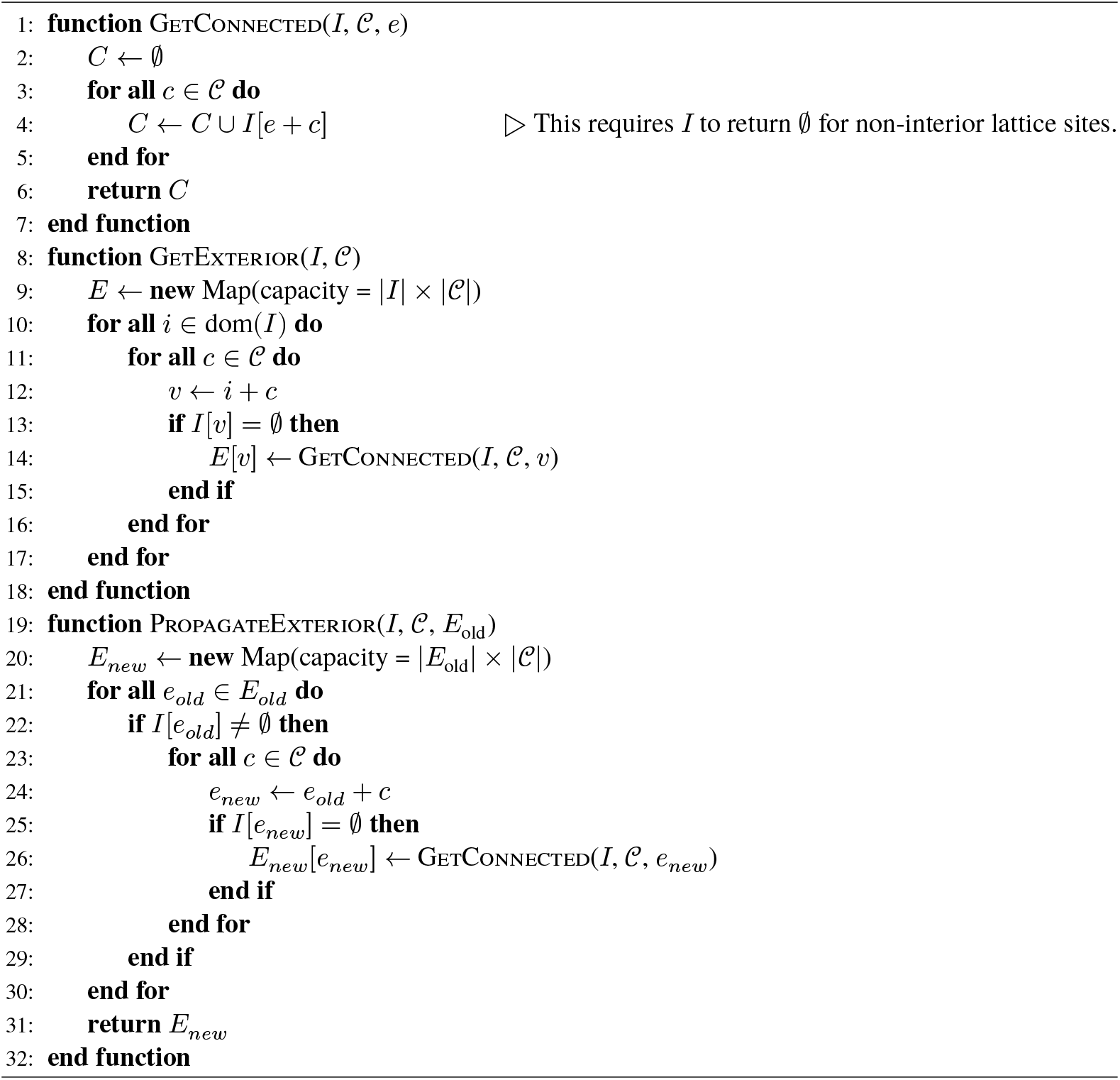

#### Algorithm B.2

Utility functions for generating interior maps, and converting between the two types. The selector function allows the user to pick the representative of each cell: for example, this may be the closest terminal to the lattice site.

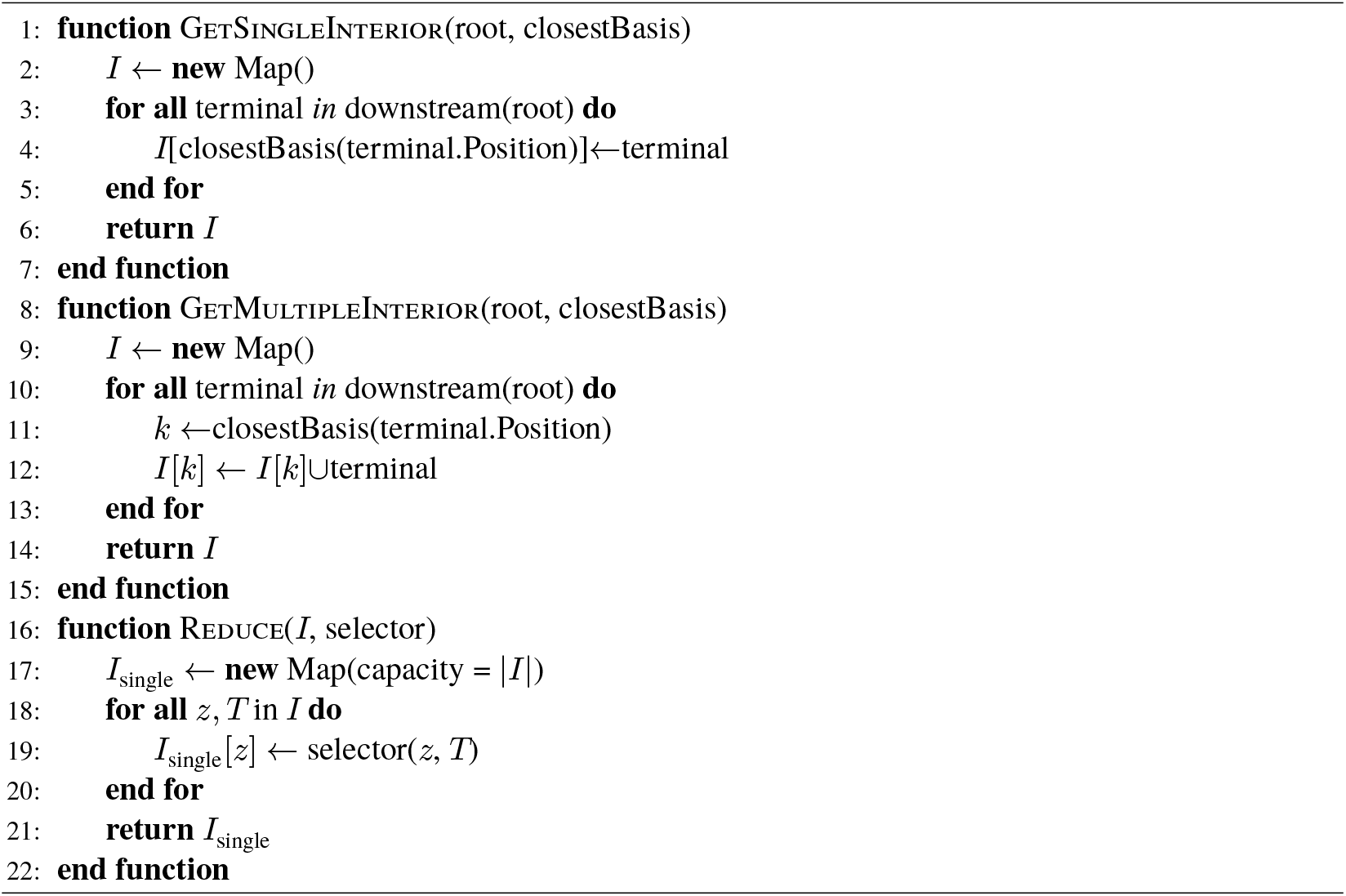

#### Algorithm B.3

The main loop. The result value determines whether more iterations may be made — if there are no more refinements to be made and *ε* = ∅, then future iterations will have no effect beyond executing the relevant callbacks.

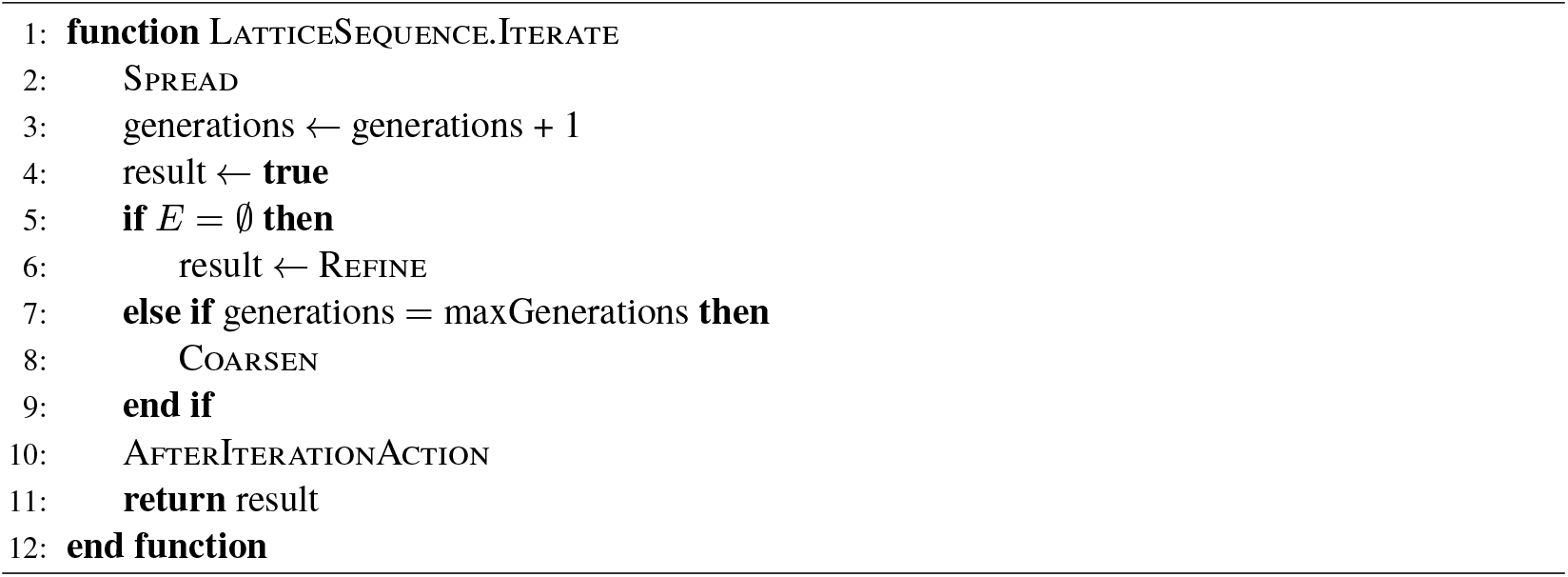

#### Algorithm B.4

The spread function. The exterior is permuted to introduce randomness into the build order, then we attempt to create a candidate terminal at each exterior site, find the most optimal existing terminal from the permissible set of connections, and bifurcate into this. Each of these functions is user specified, which creates a high set-up cost to gain customizability.

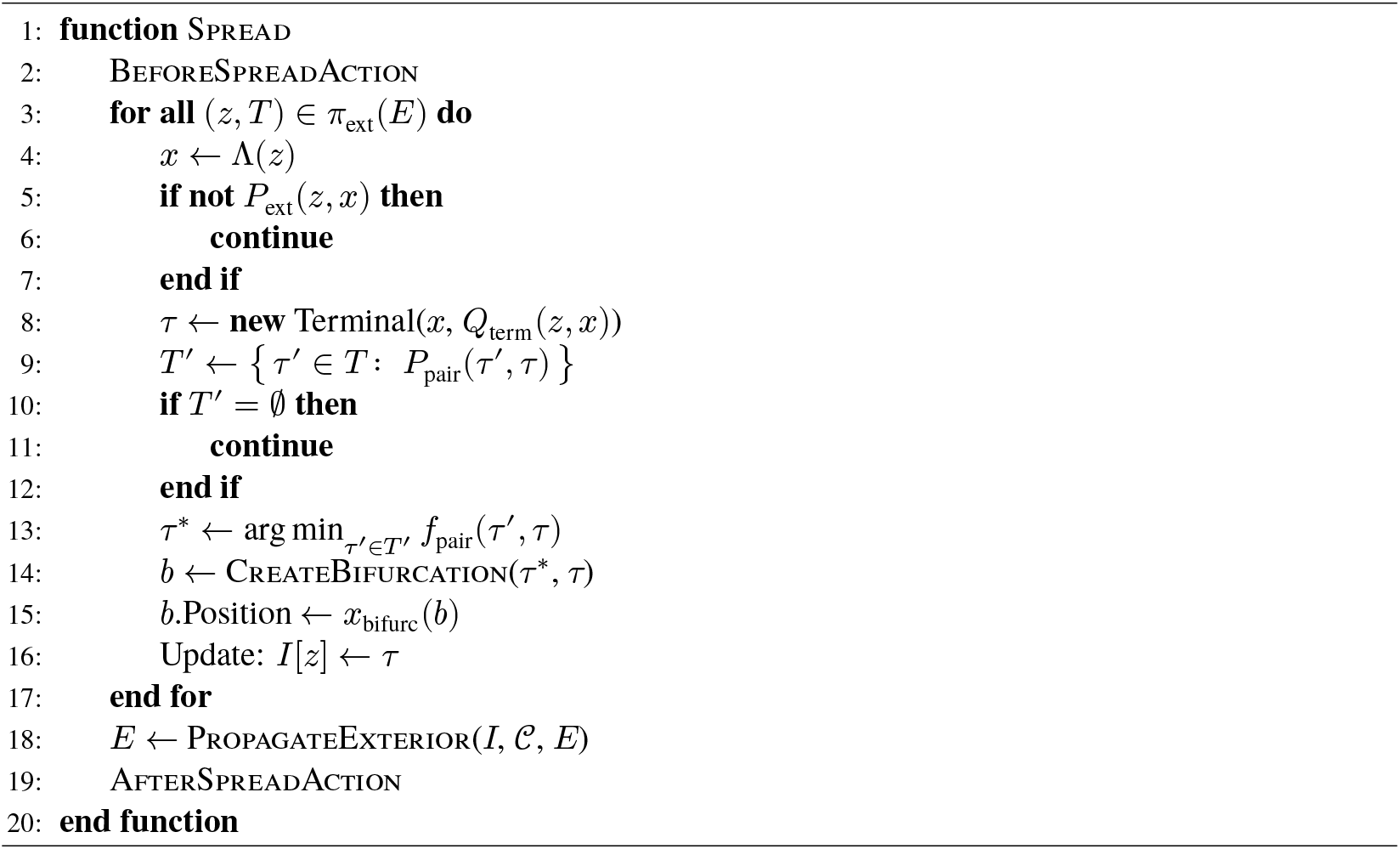

#### Algorithm B.5

The refine method of the lattice state class — this is only ever called when re-refining, so we have a guarantee that the old interior exists. The first time that a state is hit, we initialize the state instead (the refine procedure of the lattice sequence class).

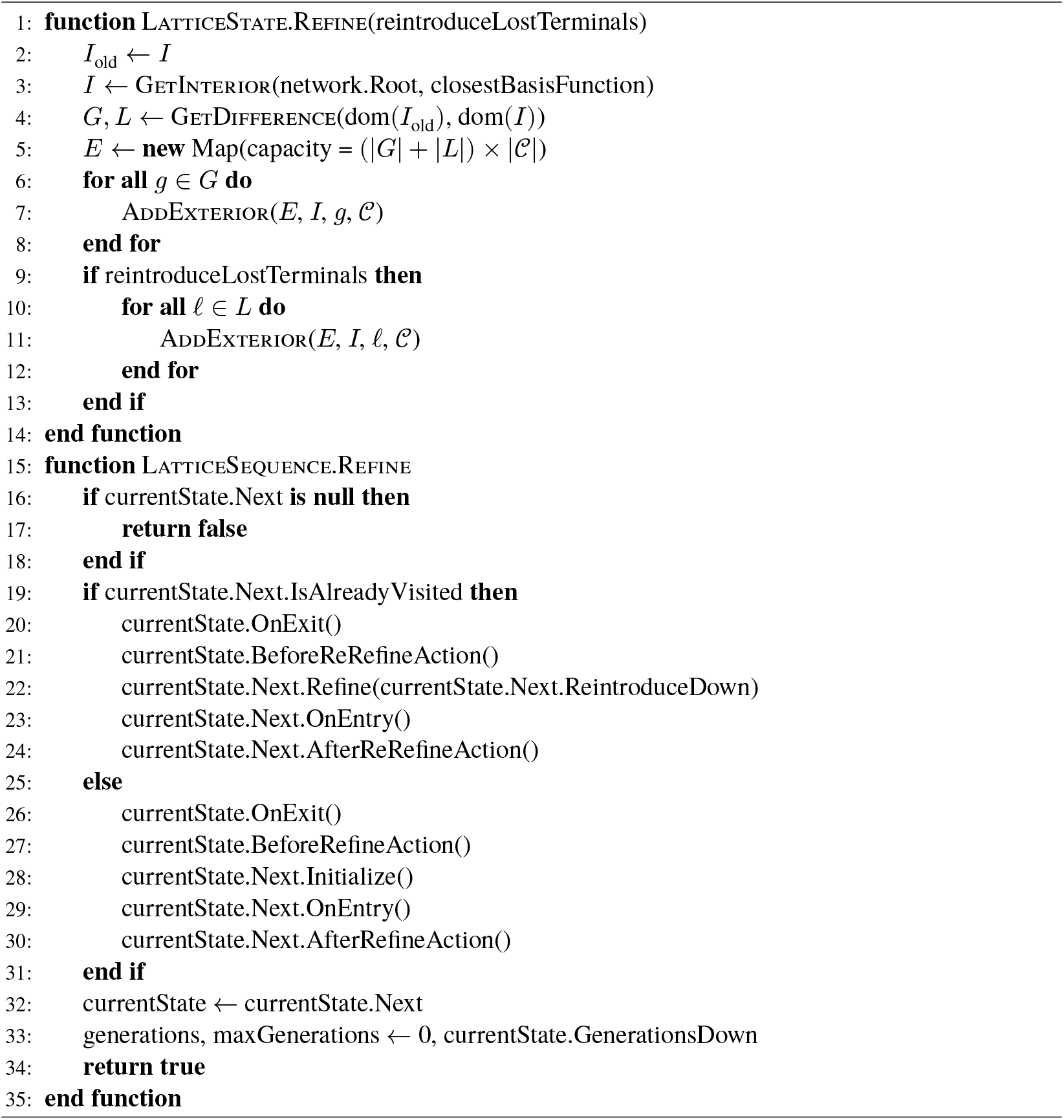

#### Algorithm B.6

The lattice state coarsening method. We reinitialize the interior and exterior from the refined state, possibly attempting to reintroduce the terminals we have lost since the last time we performed growth on this lattice. Unlike with refinement, we know that we have visited the previous lattice already, so there is only one path to take.

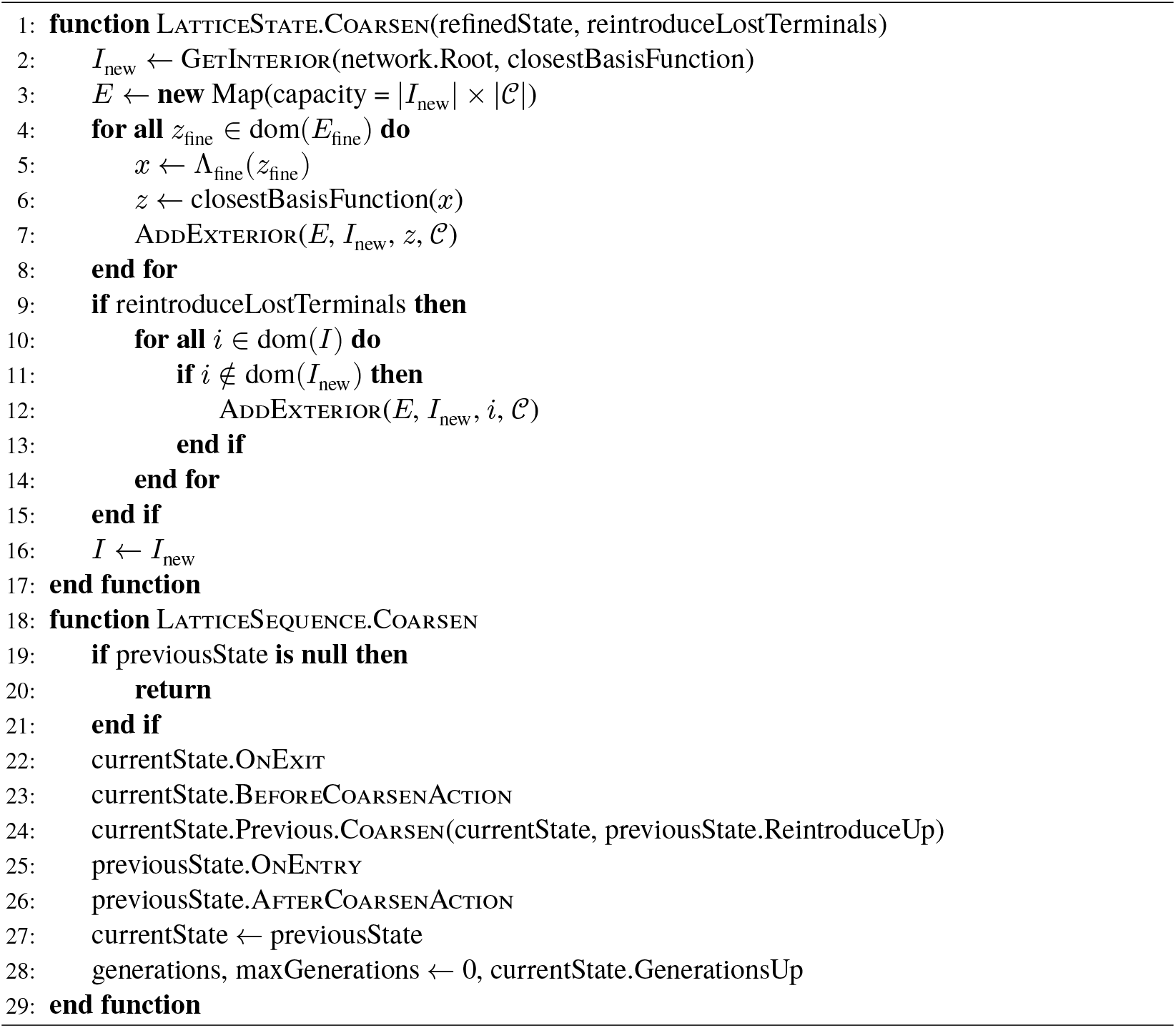

### C LSV Analysis

#### C.1 Expected computational scaling

Due to the divide-and-conquer nature of the lattice refinement process, we expect Θ (*N* log *N*) scaling for *N* terminal vessels in the final configuration. As we do not directly specify a terminal set but rather a minimum lattice stride (typically of the order of the critical cell-channel distance 𝓁^∗^), we expect *N* = Θ (|Ω|/(𝓁^∗^)^3^), which we may insert into the above. In the first analysis, we assume “typical usage”: i.e., when we increase the volume of tissue or decrease the critical length, we respond to this by extending the sequence of lattices through some (nearly) constant scaling factor. Pathological cases arising from algorithm configuration such as refining through a sequence that converges slowly to some length scale are not considered here; however, input pathologies such as the case where thin domain geometry creates a case where the early lattices cannot meaningfully contribute are considered in C.4.

##### C.1.1 Interior

For interior lattice sites, the choice of lattice sequence will determine the scaling, under the assumption that there are no unvisited interior points at the point of refinement. By Minkowski’s characterization of a lattice [75], there exists a constant *K* such that the number of points in a ball of radius *R* centered at the origin |*B*_*R*_(0) ∩ Λ| ≤ *K*(*R*^*d*^ + 1) for dimension *d*. Letting *R* be the largest distance for the previous lattice such that a *B*_*R*_(*y*) exists for some *y* with empty intersection with Λ, we have an upper bound on the number of spreading iterations for the interior sites of this lattice to be visited in the worst case; in practice, far fewer iterations are needed as the newly created vessels become involved in spreading.

With the number of iterations at each lattice being bounded by the largest value of 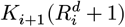 for sequential lattice pairs (*i, i* + 1), the overall computational effort required will depend on the cost of each spreading iteration (dependent on |*ε*|) and the total number of lattices. For the interior it is trivial that |*ε*| ≤ |*𝒞*|*N* for each lattice and we may take the largest of the |*𝒞*_*i*_| for this term; finally, the total number of lattices in the asymptotic scaling case depends on the minimum refinement ratio between lattice pairs. For the typical refinement sequence used in this analysis, we have Θ (log *N*) lattices and Θ (*N*) effort at each lattice.

Thus, for typical usage, we expect to see an asymptotic cost of vascularizing the interior of Θ (*N* log *N*). In practice, we are interested in relatively low *N*, and we may see very few refinements for sequences with large refinement factors.

##### C.1.2 Boundary

The overall asymptotic scaling of LSV will therefore be determined by its behavior near the boundary of the domain, in particular points which do not admit interior spheres^1^. Intuitively, the rate at which we “discover” space near the boundary as we reduce the length scale at which we spread into it will determine the asymptotic scaling, and only if this is constant will we achieve *N* log *N* scaling.

The equations to be solved can be found from a geometric construction considering the placement of an interior ball of radius *r* defined by the lattice connection pattern and its maximum distance *h* to a boundary point: the form for Lipschitz domains with modulus of continuity *K* yields a simple result that

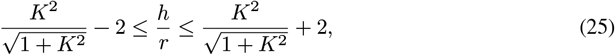

i.e., any triangulated domain will achieve *t* = Θ (*N* log *N*). This is not the case for *α*-Hölder continuous domains with *α* < 1, in which relatively more “spreadable boundary” is discovered as the refinement progresses, and this should be kept in mind when Ω is defined implicitly, as domains created by constructive solid geometry of even a few simple shapes (e.g. exclusion of cylinders touching tangentially in a containing domain) could trigger pathological behavior.

#### C.2 Bottlenecks

One of the theoretical benefits of LSV as a general-purpose algorithm for designing vasculature is that it can pass through bottlenecks in a domain, and that coarsening allows hierarchy to be developed once this has been passed (Figure C.1). Starting by defining the boundary *ε*-shell as

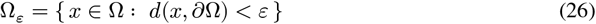

we then define the bottleneck radius as:

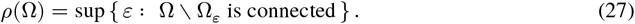

To successfully vascularize the domain beyond this, the user must first ensure that the largest vessel radius is less than *ρ*(Ω). Given a desired lattice basis and connection pattern, we have a radius proportional to ‖Λ‖_2_ in which a guaranteed connection pair exists that does not leave a ball of this radius; hence, we can guarantee passing a bottleneck if one of the lattices in the sequence has a small enough connection radius, and this is again the responsibility of the user.

The feasibility of this approach is predicated on the bottleneck being broad enough to accommodate an appropriately-sized vessel for the flow rate required to supply the tissue on the other side of the bottleneck. Furthermore, optimization and collision resolution routines may move the vessels of this feasible network to intersect the boundary near the bottleneck, requiring constant testing and possibly causing the same pinning issues that were prevalent with existing CCO approaches for interpenetrating networks, so users should consider whether bottlenecked domains are best handled as distinct components to be vascularized separately (perhaps providing a predefined major vessel path between these) before embarking on a potentially expensive growth process.

#### C.3 Meeting a critical distance

Thus far, scaling analysis has been asymptotic, rather than considering the minimum amount of computational expense required to meet manufacturing constraints. Again, the boundary will determine the minimum lattice spacing required to satisfy the critical cell-channel distance. For triangulated domains, we have already established the estimate 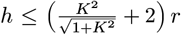, therefore the required lattice spacing (as *r* ∝ ‖Λ‖_2_) depends on 𝓁^∗^ and the sharpest angles of the triangulation.

We note that in practice, the combination of smallest manufacturable feature sizes, desired cell-channel distances and practicalities of manufacturing (either *in situ* by ablation or fugitive ink processes, or by mounting 3D-printed templates) will render the question of growing vessels into long and thin or coiled (e.g., “pigtail”) boundary portions irrelevant, even though these would otherwise provide a pathological case in which the naïve determination of lattice spacing would lead to necrosis at the tips of these portions. Thus, for bulk tissue engineering, the domains encountered are naturally constrained to be non-pathological.

#### C.4 Pathological cases

For domains consisting of thin shells with curvature, it may not be possible to grow on a coarse lattice and attain the required hierarchy for good performance. In this case, we expect that the factor of log *N* associated with refinement will degrade to 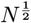 (associated with spreading over the entire length, for the most pathological domains which are thin in two dimensions this will degrade further to *N*) in the range of *N* corresponding to ‖Λ‖ which are larger than the maximum admissible stride, but asymptotic performance for *N* → ∞ will remain unchanged provided that the shell has finite thickness (for our current use-cases, we are not going to pass into this region). Such cases may arise when designing templates for tube walls, myocardium or the choroid of the eye, and the optimal approach in this case may be to develop simple analytical descriptions of the surfaces and supplement the optimization with a penalty term quantifying the violation, or to grow in more favorable shapes and map into the final structure, accepting a small deviation (see [38] for the application of this approach to the retina).

**Figure C.1:**
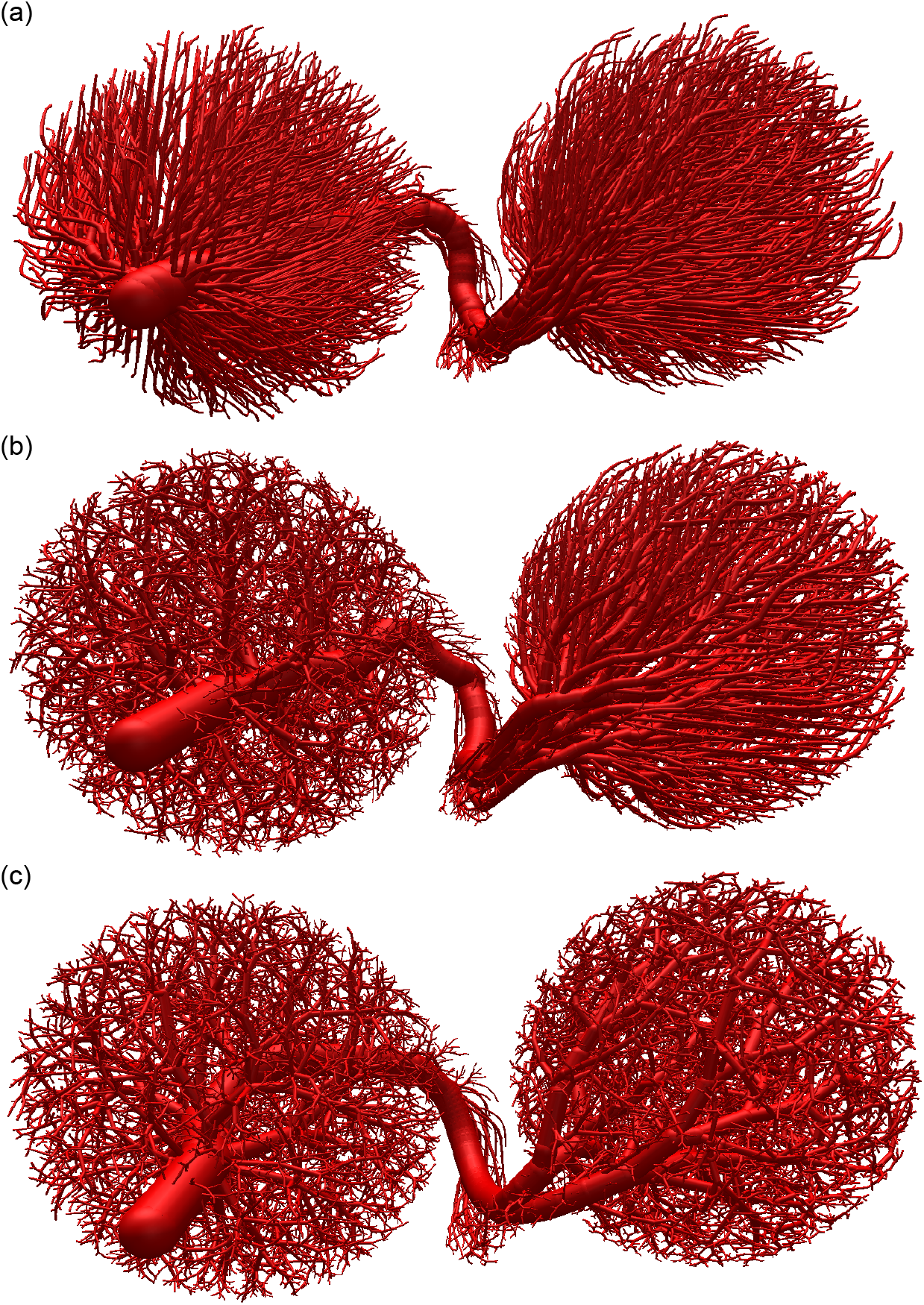
The motivation for LSV: (a) passing a bottleneck (b) refining in the first ball (c) coarsening after passing a bottleneck.

**Figure D.1:**
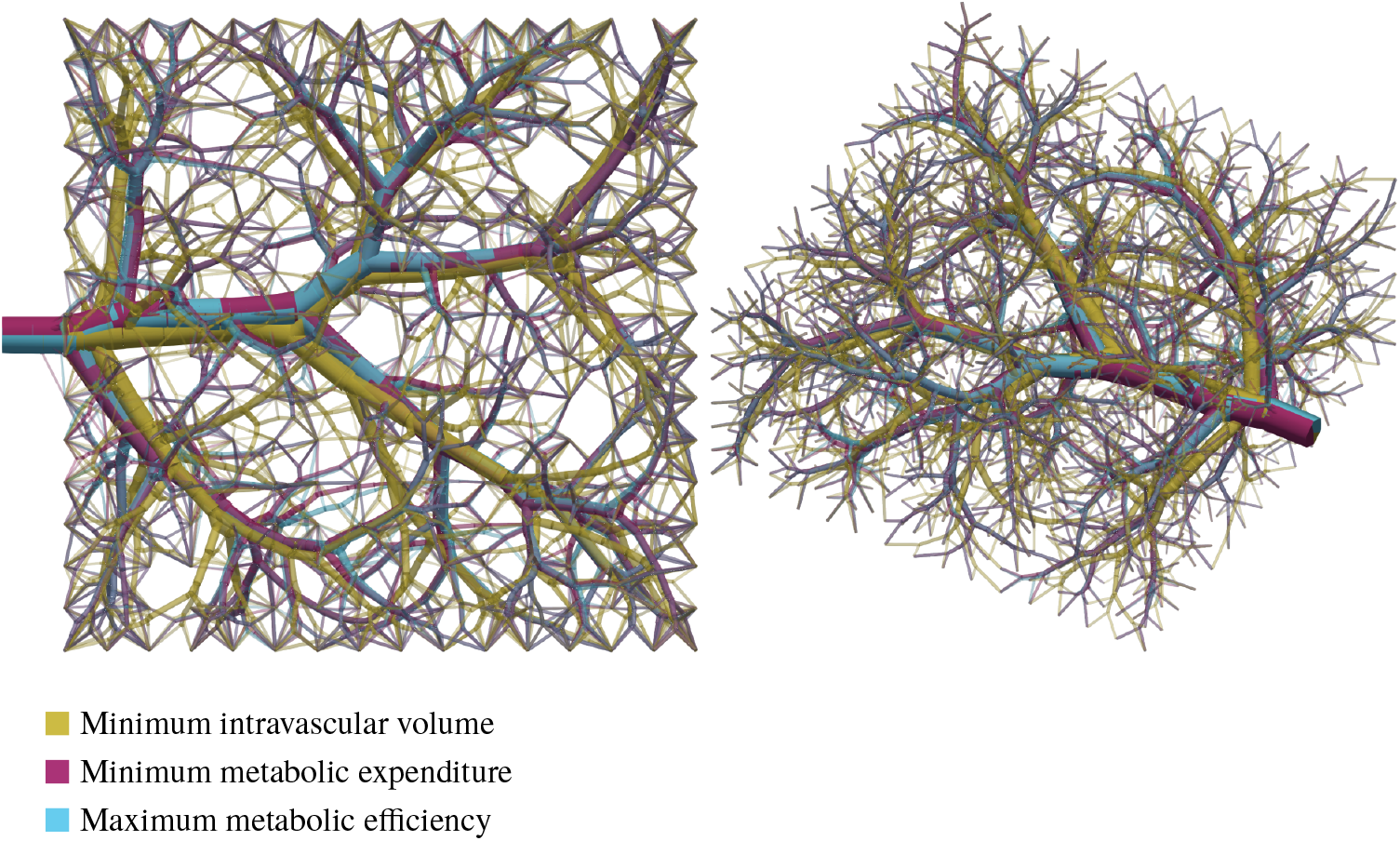
Three vascular networks generated from the same seed and optimized for different costs: minimum intravascular volume, minimum metabol1ic expenditure, and maximum metabolic efficiency. Despite the differing optimization costs, the resulting network structures are remarkably similar.

### D Optimization

#### D.1 Differences between volume-dominated costs

Figure D.1 shows three networks arising from the same seed, where the costs are the minimum intravascular volume (i.e., maximum useful tissue), the minimum metabolic expenditure (i.e., the cheapest construct to maintain) and the maximum metabolic efficiency (the ratio of the two preceding costs, representing maximizing the “output” per unit energy expenditure), respectively, where the metabolic expenditure per unit blood volume was taken from [76]. It can be seen that for all three costs, the resulting network structure is extremely similar, suggesting that the metabolic expenditure of maintaining blood volume reported in the literature is dominating.

#### D.2 Geometric optimization

In [30] we showed that networks with lower mean cost and lower variance could be generated using CCO by replacing “incremental optimization”, in which newly created bifurcations are moved in isolation to minimize the network cost, with “batch optimization”, in which the growth process is interrupted and all nodes moved simultaneously. We also showed a simple approximation of the gradient direction using only local data, which was shown to align well with the true gradient, and that placing the initial bifurcation at the flow-weighted arithmetic mean of the surrounding nodes lead to a much lower cost by better approximating the optimal location. Other work has shown that the approach of optimizing the entire tree at once yields better results than incremental approaches [20, 31]. In this work we formulate a Θ (*n*) exact gradient computation which relies solely on the isobaric terminals assumption, further improving optimization results.

##### D.2.1 Hierarchical properties

Efficient update rules for reduced resistances and radii fractions have been a core part of CCO methods since the original implementation [12], recognizing that changing a single vessel need only propagate the changes back up the direct path upstream to the source rather than recalculating all vessels. This work builds on the “hierarchical costs” concept of of our previous work [30], which allows efficient update of polynomial costs over the set of branches *ℬ* of the form 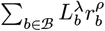 as considered in [54] by the *effective length* relationships at terminals and bifurcations:

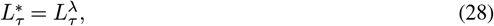

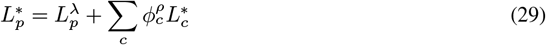

which allows the cost at the root to be expressed as 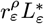.

##### D.2.2 Gradient chain

We first partition our properties into two sets: *(child-) derived* (*A*), which include *R*^∗^, *Q, L*^∗^; and *(from-parent-) assigned* (*B*), which includes *ϕ*. We then have an *assignment function B*_*c*_ = *g*(*A*_*c*_), and then the parent *derivation function A*_π_ = *f*(*A*_*c*_, *B*_*c*_), where *g* : ℝ^*k*×*n*^ → ℝ^*k*×*m*^ and *f* : ℝ^*k*×*n*^ × ℝ^*k*×*m*^ → ℝ^*n*^ for *n* derived properties and *m* assigned properties with *k* children. In this implementation, the only assigned property considered is *ϕ*, the radius fractions between parent and child branches.

Now consider the impact on a root property 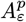 from a downstream property at an arbitrary address 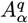 — this is how we will calculate gradients of hierarchical costs. For any node:

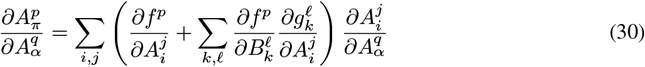

where the subscripts *i, k* span the children and *j*, 𝓁 the properties. By the hierarchical properties principle, 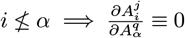 . Thus, at a parent node π, only one of the child nodes will have non-zero terms for the address *α*. Let this on-chain child be denoted by the index *c*:

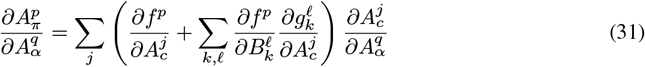

we see that this is a linear mapping of an *n* × *n* matrix of derivatives from π to *c* — so for a single-pass calculation we must simply construct this matrix.

For practical purposes, we need not maintain the entire *n* × *n* state. Setting

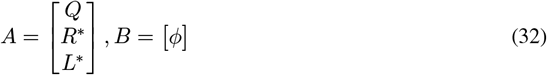

we then have:

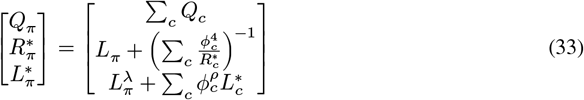

where we use a partially non-dimensionalized form of *R*^∗^ to remove constant viscosity and geometric scaling factors, and for a generalized splitting rule:

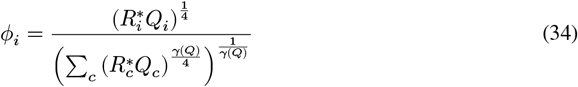

and the chain matrix is therefore:

**Table D.1:** The number of intermediate variables required to be passed downstream to calculate the hierarchical gradients of a root property with respect to both position and flow rate at a node. The effective length gradients are tabulated for *n* instances of (*λ, ρ*) pairs, as they have a dependency on the underlying *R*^∗^ and *Q* gradients which need not be recalculated.

| | $Q_\varepsilon$ | $R_\varepsilon^*$ | $n \times L_\varepsilon^*$ |
| --- | --- | --- | --- |
| $x_\alpha$ | - | 1 | $1 + 2n$ |
| $Q_\alpha$ | 0 | 2 | $2 + 3n$ |

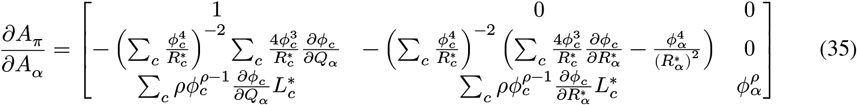

Thus, for calculating gradients with respect to the node/branch properties of interest we require the number of intermediate variables as specified in Table D.1. Note that the gradient 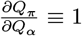 need not be passed, and that multiple polynomial costs do not interact with each other. In practical usage, the gradient for each polynomial cost may be calculated independently, re-using cached values of gradients with respect to *Q, R*^∗^. For node positions, we then calculate the gradients of *L* for all attached branches, and then use the relevant properties (*R*^∗^, *L*^∗^) to get the root gradient.

##### D.2.3 Gradients of radius fractions

In the computation of gradients for *R*^∗^ and *L*^∗^, we see the terms 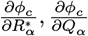. For Murray’s law, we may write 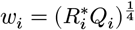 and we then have

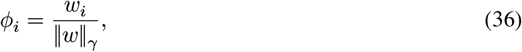

giving the direct the gradient with respect to the child *i*:

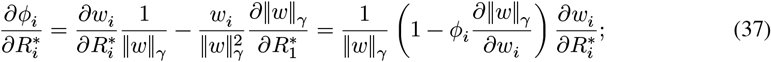

and cross-terms:

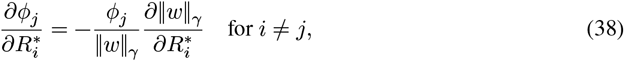

Where

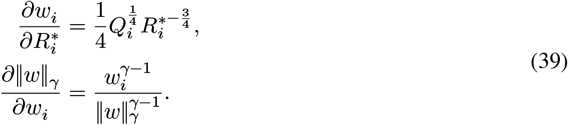

Combining these together, for constant *γ* we have:

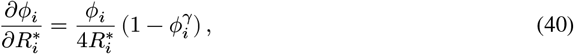

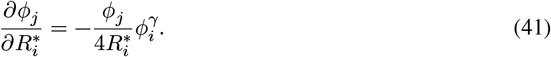

For the variable Murray’s laws that have been implemented, where the exponent may depend on the flow, we have extra terms arising from the fact that

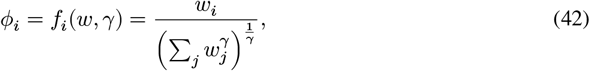

where both *w* and *γ* are functions of *Q*. As the splitting law is defined in terms of the parent flow, we know that 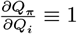, and we have

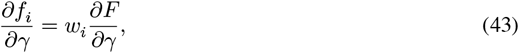

where

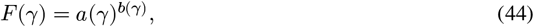

With 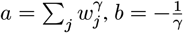 . We therefore have

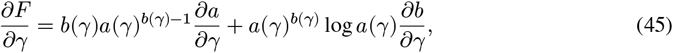

with 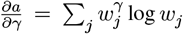 and 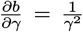. We may evaluate 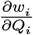 in the same fashion as with *R*^∗^, and reassemble the terms to get

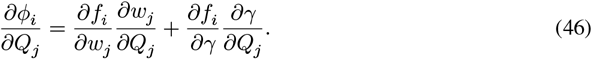

### E Simulated Annealing

A simulated annealing approach for topological optimization of networks has been developed based on the work of Keelan *et al*. [20]. We note from our own work [30] that the impact of “deep” nodes on the overall network cost is low; previous studies have found that the structure of the root vessels has a great impact on the final cost when the network is grown further [14, 54, 57]. Therefore, we conclude that the approach of [20], in which the network is completely constructed before simulated annealing begins, is extremely wasteful due to the rapid growth of the number of topologies, and the fact that terminal-scale vessels outnumber the root vessels. Instead, we propose that simulated annealing be employed at an early stage, where all terminal vessels represent a volume that will eventually contain many other vessels. The network can then be grown from this starting point. Further, we should not start from a truly random starting network, but use a biomimetic growth procedure to generate networks which are known to have acceptable costs. This reduces both the number of topologies that must be considered and the expected perturbation distance between start and end configurations, requiring fewer iterations to be confident of reaching a good solution, as well as ensuring all changes are meaningful. Work by Jessen *et al*. [31] showed this hypothesis to be correct, employing simulated annealing between CCO growth at *N* = 500 for 10^5^ iterations and achieving a reduction in volume of up to 6% compared to geometric optimization alone when applied to a standard test case specified by Karch [13]. They used a lower starting temperature and more rapid annealing schedule than [20], justifying this by their more optimal starting configuration compared to the arbitrary initial network.

Whereas Keelan *et al*. introduce perturbations by moving a node or swapping a bifurcation, we replace the node movement step with gradient descent applied between topological changes, thereby reducing the overall number of iterations required for convergence. This is the same top-level approach as [31] although their geometric optimization approach differs in that it is performed by an NLP solver, as well as the problem they are solving: minimizing volume with node locations/pressures and branch lengths/radii in the solution vector and equality constraints of Murray’s law and length/pressure consistency.

Further, we assume that our initial state, having been generated by a biologically-inspired process, will be somewhat near-optimal, and that we do not need to make large changes to the topology. Therefore, we instead generate perturbations by random promotions of nodes, reducing the number of candidate actions each iteration to Θ (*n*) as opposed to Θ (*n*^2^) bifurcation moves — if our network is nearly optimal, this will increase the likelihood of beneficial moves being selected as it prevents distant vessels from being attached to each other, reducing the number of iterations required for convergence at the cost of reduced capacity to explore the search space.

**Figure E.1:**
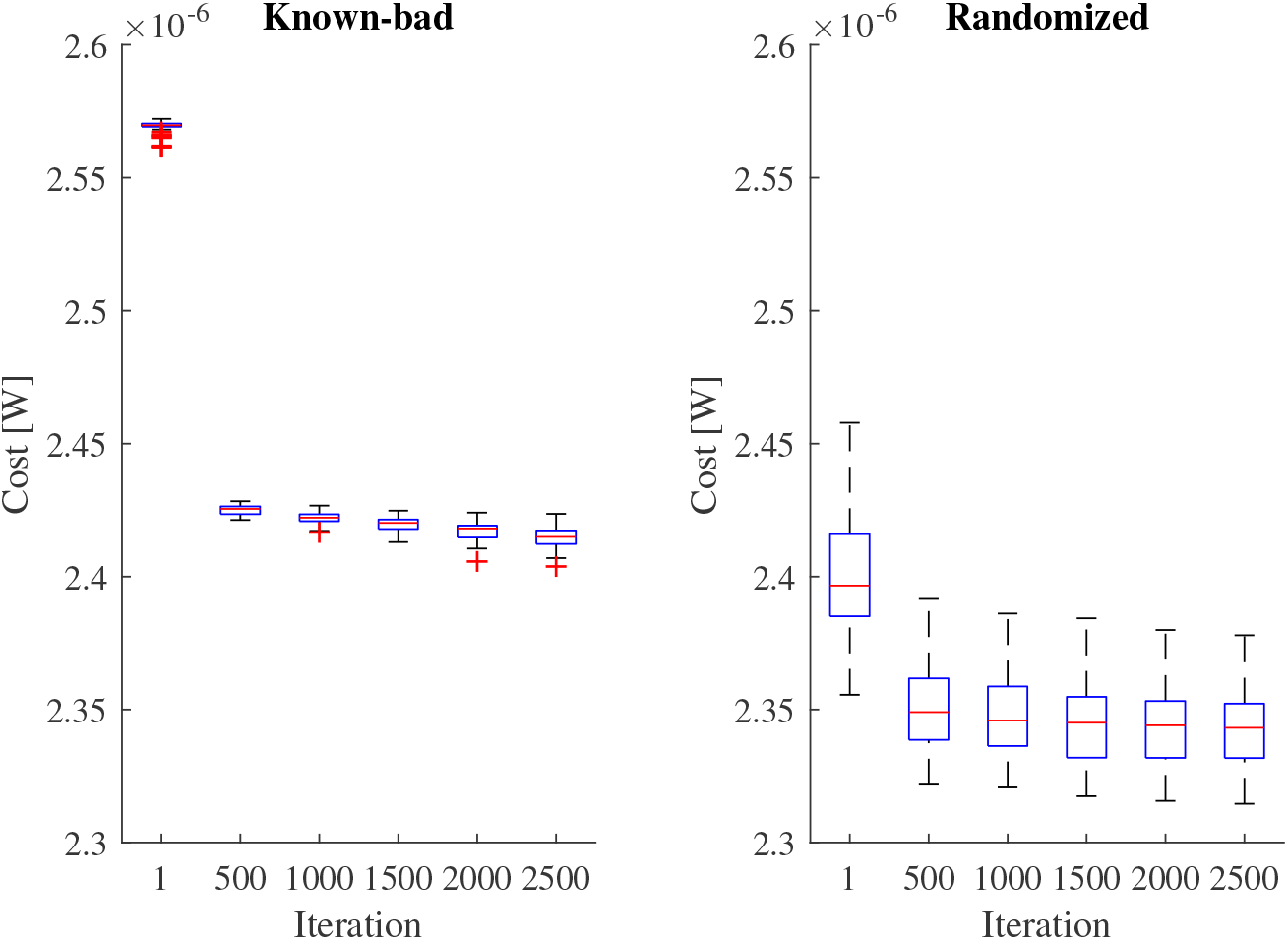
Distribution of costs for 100 instances of a 6×6×6 cube network grown by ACCO. Left: a known bad starting network, grown without optimization and shuffling. Right: a non-optimized starting state, but with randomized growth order.

It can be seen that the initial topology has a clear impact on the result, due to the rapid growth of the number of topologies with terminal count making the search space infeasibly large. Figure E.1 shows the evolution of costs for a known-bad 6×6×6 terminal network, grown using ACCO without randomization in the terminal order, as well as the results of the same procedure but starting from a network where the terminal construction order was randomized. Better results are achieved by using strategies which avoid creating topologically non-optimal networks (e.g., biologically-inspired procedures): the mean cost is 3.0% lower; and, when compared to the minimum cost observed when starting from the known-bad network, the randomized starting point leads to networks with costs 1.1–3.7% lower.

Figures E.2 and E.3 show this approach applied to the same setup as in [20], with a 10 cm square fed by an 8×8 terminal square, with *μ* = 3.6 × 10^−3^ Pa⋅s and a total flow rate of 4.16 × 10^3^ mm^3^s^−1^. It can be seen that with only 5000 iterations, we can converge to similar structures and costs as they achieved in 5 × 10^9^, a factor of 10^6^ fewer iterations.

### F Higher order splits with variable Murray’s law

One of the advantages of constant Murray’s exponent is that we are able to simulate higher-order splits (*N*-furcations) using only bifurcations, which we show here. Consider a set of downstream nodes drawing from an index set *I* with flow rate and reduced resistance *Q*_*i*_, 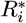, let the connectivity be given by a tree topology 𝒯 ⊂ 𝒫 (*I*) and let them have separation low enough (i.e., that the intermediate branch lengths are effectively zero) such that the reduced resistance of the intermediate branches is given by:

**Figure E.2:**
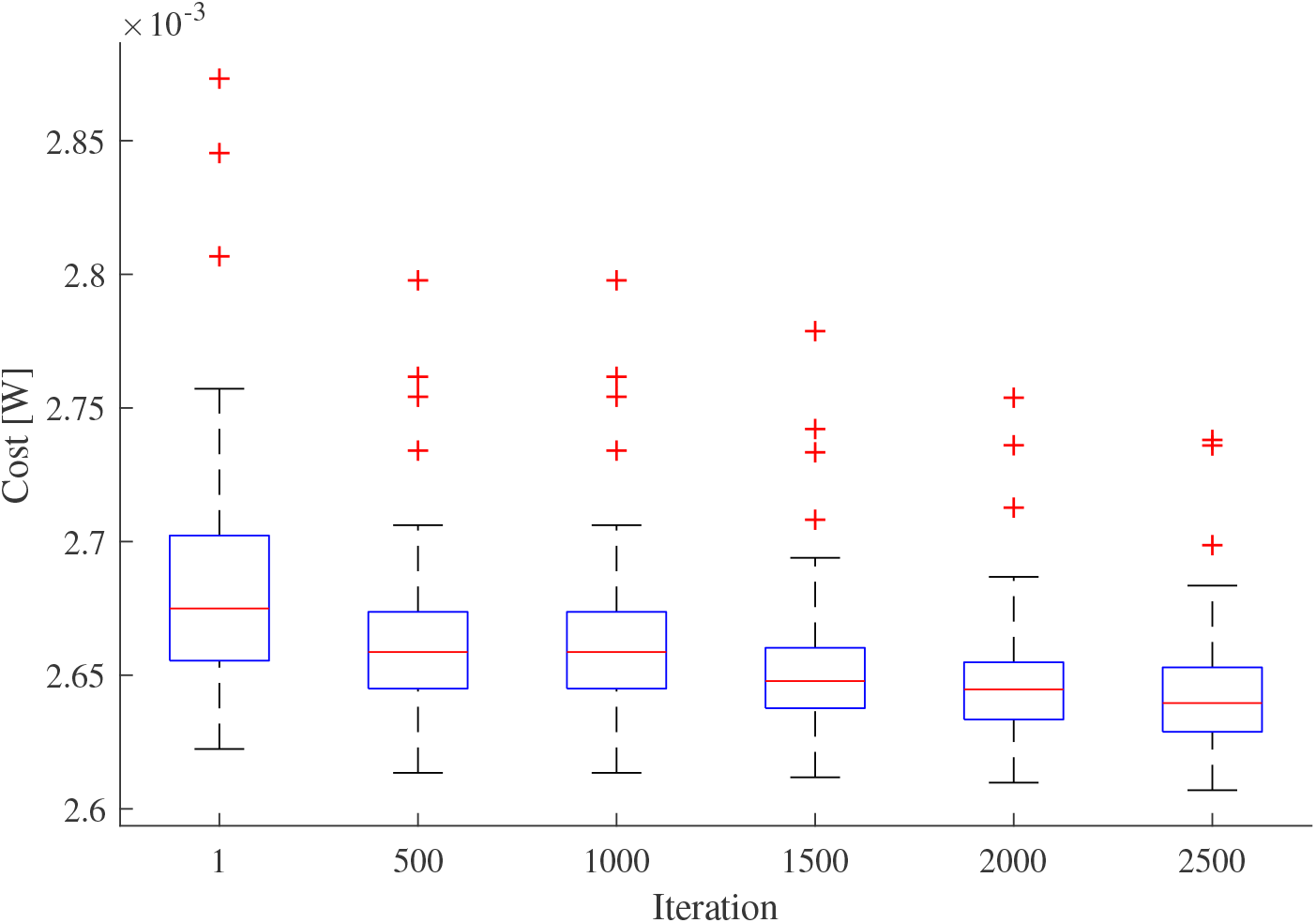
Cost of 100 instances of the same test configuration used to generate Figure 9 in [20], after a given number of iterations.

**Figure E.3:**
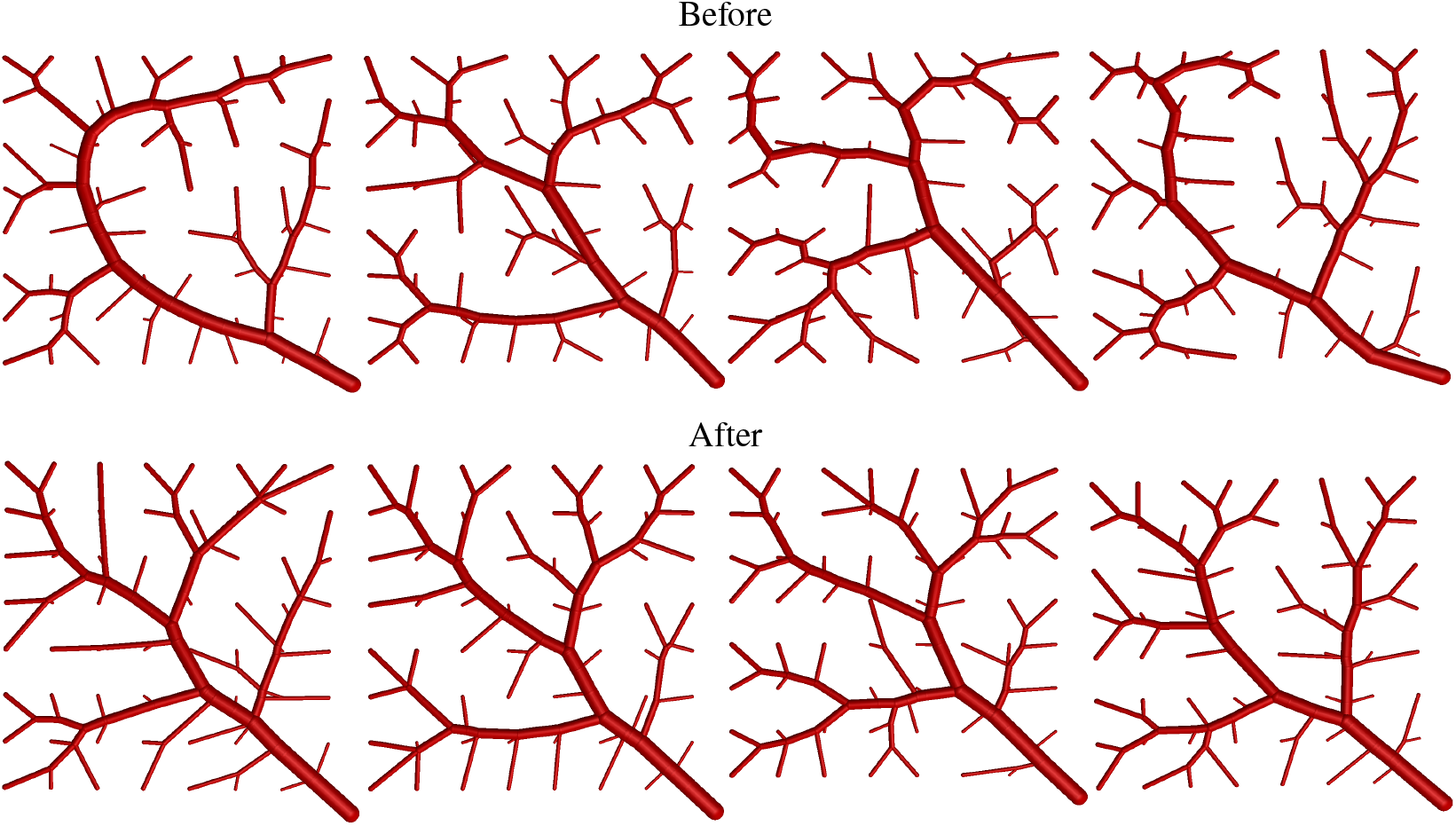
Examples of the same configuration as tested in Figure 10 of [20] after 5000 iterations of the modified simulated annealing scheme.

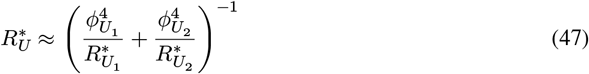

for *U*_1_ ∪ *U*_2_ = *U, U*_1_ ∩ *U*_2_ = ∅ and *U* ∈ 𝒯. Now, consider a variable Murray’s law: we have, for *V* = *V*_1_ ∪ (*U*_1_ ∪ *U*_2_), with *V*_1_, *U*_1_, *U*_2_ disjoint, we have the intermediate fractions:

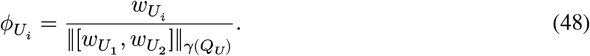

We then pass these upstream as

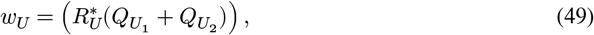

and we insert the radii ratio to yield

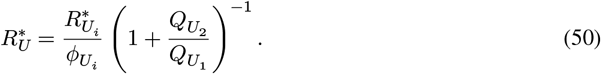

We therefore have

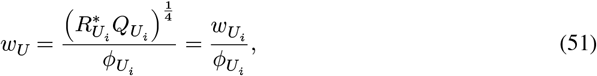

and we see that any choice of splitting function will lead to overall fractions that lie on a line *ϕ*_*i*_ ∝ *w*_*i*_, as long as it constrains each pair of intermediate pseudo-vessels to lie on a line 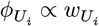.

For a Murray’s law type of splitting function, we have the relationship at an intermediate node due to (48):

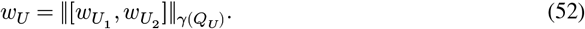

and therefore at the node *V* we have

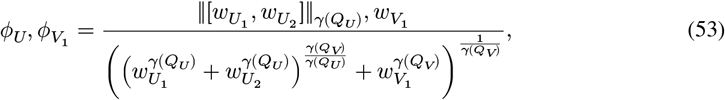

and we may chain the relationship for intermediate fractions to find that overall, we have

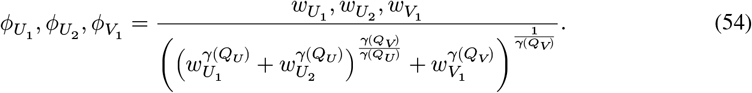

The form of this is as expected, as the isobaric terminals assumption means that we are looking for a scaling factor to scale the *w*_*i*_ of the endpoints. If *γ*(*Q*) is constant then the terms *γ*(*Q*_*V*_)/*γ*(*Q*_*U*_) = 1, and the entire function becomes *L*^*γ*^ normalization, enabling the simulation of higher-order splits using only bifurcations in an order-independent way. We note that higher-order splits are rare in real vasculature, but can be a useful intermediate step in optimization to enable a transition to a more optimal network topology. The full treatment of derivative calculations requires a *d* ×*d* matrix for splitting order *d*, which makes limiting the maximum splitting order imperative for good performance when using the scheme presented here, and makes schemes such as GCO impractical, as this method starts with a single splitting node branching directly to all terminals.

### G Radius smoothing reduces optimality

Another interesting property is the concept of branch profiles: where radius is non-constant along the branch. This could be used to reduce the impact of sudden changes in radius (such as step changes in wall stress or deviation from the pipe flow assumptions at bifurcations), as well as being more biomimetic, allowing a smooth transition between vessels. This is represented using a scaling factor σ > 0 as a function of position along the branch length and impacts derived properties as such:

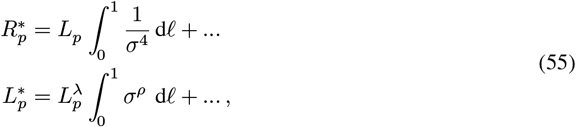

where 𝓁 represents the length fraction along the branch. Note that in the case where *λ* ≠ 1, it is difficult to assign meaning to the alternative formulation of the sum over branch segments 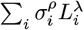, as a fragmentation of a branch with σ ≡ 1 would have a different cost: the definition in (55) does not have this issue.

Whilst we could in theory provide any σ, we would in practice restrict ourselves to those satisfying:

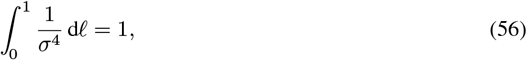

else the reduced resistance of the branch would be modified as though the branch were a different length, and in order to supply the correct flow rate we would end up assigning a correspondingly different radius. Considering the impact on optimality, we inspect the modified functional:

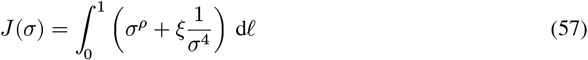

where *ξ* is a Lagrange multiplier, yielding the Euler-Lagrange equation

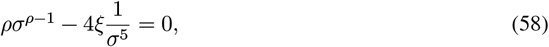

with solution σ ≡ 1 when 4*ξ* = *ρ*, the unity scaling. Noting that the impact on effective length is a multiplication by 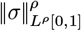, we appeal to the convexity of norms to conclude that the application of any non-unity profile under the constraint (56) will decrease the optimality of polynomial costs for *ρ* ≥ 1, whilst leaving pump work and polynomial costs with *ρ* = 0 untouched. We therefore consider branch profiles to be a post-processing step, to be executed alongside the final round of constraint enforcement — we cannot apply a profile after constraint enforcement has run as it must expand the branch in at least one segment, which may lead to intersections. For our demonstration, we have used Laplacian mesh smoothing with flow rate weighting as a simple approach to minimize radius jumps along paths delivering large flow rates.

### H Liver terminal lattice

The functional structure of the liver arises from its lobular tiling of portal triad (portal vein, hepatic artery, biliary tree) and venous penetrations. There are 2 portal columns for each venous column, arranged in a hexagonal tiling as shown in Figure 3(c). It can be seen that if the basis vectors are chosen as shown, calling them *a*_1_, *a*_2_ arbitrarily, then the hepatic vein sites occupy the sublattice with basis:

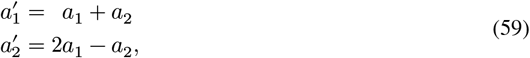

and we wish for alternating planes of portal and venous terminal vessels, between which the functional structure may then be constructed between the sheets as penetrating columns with interconnecting capillary vessels. This structure can be replicated by supplying a hexagonal prismatic lattice as the final stage which looks like, up to rotation and offset:

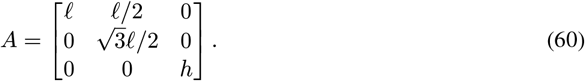

We then grow the venous vessels on a lattice that is scaled by a factor of 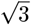 in the XY-plane and rotated by π/6, and the portal triad can be grown with an exterior predicate to knock out terminal sites occupied by the venous lattice:

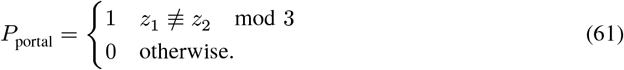

The connectivity is restricted to the 6 in-plane neighboring lattice sites, and the alternating sheet structure is created through a *z*-offset.

## Footnotes

1 i.e., there does not exist a ball *B*_*r*_(*y*) ⊂ Ω for the point *x*_0_ ∈ *∂*Ω such that 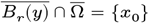. Regions which admit interior spheres will behave the same as interior points.

## Notes

### Competing Interest Statement

The authors have declared no competing interest.

